# Functional characterization of a Con7-related transcription factor in *Coprinopsis cinerea* indicates evolutionary conservation of morphogenetic roles

**DOI:** 10.1101/2025.09.11.675558

**Authors:** Hongli Wu, Zsolt Merényi, Máté Virágh, Xiao-Bin Liu, Botond Hegedüs, Zhihao Hou, Edit Ábrahám, Anett Fürtön, Zsolt Kristóffy, Zoltán Lipinszki, László G. Nagy

## Abstract

Fruiting bodies of mushroom-forming fungi (Agaricomycetes) exhibit the highest degree of multicellular complexity in fungi, yet the molecular underpinnings of their developmental programs remain incompletely understood. Here, we characterize *gcd1*, a gene encoding a transcription factor in the Con7 subfamily of C2H2-type zinc finger proteins. This subfamily has previously been implicated in pathogenic morphogenesis in Ascomycota, but their role in Agaricomycetes has not previously been addressed. In *Coprinopsis cinerea*, CRISPR/Cas9-mediated deletion of *gcd1* resulted in strains with severely impaired fruiting body morphogenesis, with malformed cap, stipe, and gill tissues. *Gcd1* deletion strains lacked universal veil, resembling species with open (gymnocarpous) development. We find that GCD1/Con7 homologs are widely distributed in most Dikarya species and are mostly encoded by a single gene in each species’ genome. Transcriptome analyses identified several misregulated genes in the Δ*gcd1* mutant, which pinpoint potential mechanisms underlying its developmental defects as well as provided insights into the morphogenesis of mushroom fruiting bodies. These findings establish GCD1 as a key regulator of multicellular development in *C. cinerea* and broaden the known functions of Con7-like transcription factors to include fruiting body morphogenesis in Agaricomycetes. Overall, our results and the morphogenetic role of Con7-like transcription factors of Ascomycota suggest functional conservation over half a billion years of evolution.

## Introduction

Fruiting bodies of mushroom-forming fungi (Agaricomycetes) represent some of the most complex structures produced by fungi. Their development is governed by a genetically encoded developmental program that involves intricate regulatory mechanisms, most of which are yet to be elucidated. This leaves the basic principles of mushroom development and evolutionary transitions between morphotypes poorly understood. A key challenge in fungal developmental biology is uncovering developmental genes and the regulatory logic that underlies the development of sexual fruiting bodies.

Recent research has made significant progress in documenting the genetics of fruiting body development and uncovered several regulatory genes, mostly transcription factors, that, in various species, regulate diverse aspects of fruiting. For example, in *Schizophyllum commune* transcription factors have been described that regulate the initiation and number of fruiting body primordia^1,2^, biomass production, or immune responses within developing mushrooms^3^. In *Coprinopsis cinerea* less than a handful of transcription factors have been characterized, which currently cover patchy aspects of fruiting body development, such as light responses^4^, cap expansion^5,6^, gill formation^7^ and spore morphogenesis^8^. Despite these advances, current knowledge is patchy in any model mushroom species and lacks key details on the differentiation process within mushroom initials.

In a typical mushroom-forming fungus, such as the model species *C. cinerea*, development proceeds from undifferentiated hyphal knots (also called aggregates or initials), through several primordium stages in which tissue types - e.g. caps, gills, hymenium and stipe are established in a protected environment, to mature fruiting bodies that sporulate^9,10^. However, there is a large diversity in how fruiting bodies develop in various species. For example, early developmental events in some Agaricales species are exposed to the external environment (called gymnocarpic or exocarpic development) while in others basic tissues are established within a primordium that is ensheathed by protective veil tissues (called angiocarpic development^11,12^. Among model species, the ink cap mushroom *C. cinerea* represents an example of angiocarpy, with both partial and universal veil covering primordia, whereas *S. commune* is a gymnocarpic species with a unique developmental process that evolved via the aggregation of simplified, open cup-like (so called cyphelloid) fruiting bodies. Gymnocarpy was shown to be ancestral in the Agaricales, whereas angiocarpy evolved several times independently and likely conferred adaptive advantage that led to higher speciation rates compared to gymnocarpic species^13^. While gymno- and angiocarpy represent two very basic developmental types, the underlying genetics is currently unknown.

The defining tissue types in gymno- and angiocarpic fruiting bodies are universal and partial veils, which, in early stages, cover the whole fruiting body and connect the cap margin to the stipe^11,14^, respectively. In mature fruiting bodies universal veils are recognized as scales, patches or dots on the cap and/or as bag-like sack at the stipe base (called volva). Partial veil is most often left behind as a ring or annulus on the stipe. The presence of veils and the shape, color and size of their remnants are important diagnostic characteristics in the taxonomy and classification of mushroom-forming fungi. They are also subject to strong evolutionary selection^14^ and may be breeding targets in commercially produced species in which sturdy fruiting bodies with a long shelf-life and delayed cap-opening are preferable.

In this study, we examined the function of a conserved, cap-expressed transcription factor, referred to as *gcd1*, in *C. cinerea* and show that it regulates primordium development and is related to gymno/angiocarpy. Deletion of *gcd1* led to defects of fruiting body differentiation, resulting in malformed cap, stipe and gills and the absence of veil tissues. Phylogenetic analysis revealed that *gcd1* is evolutionarily conserved across fungi, from unicellular yeasts to complex multicellular mushrooms. Transcriptomic analysis of our *gcd1* mutant identified biochemical pathways that might explain the observed morphogenetic abnormalities and uncover new elements of fruiting body development. These findings provide new insights into the molecular mechanisms underlying tissue formation, differentiation and developmental types in mushroom-forming fungi.

## Results and Discussions

### Identification of a cap-specific transcription factor gene and generation of deletion mutants

As data accumulates on the developmental transcriptomics of Agaricomycetes species^10,15^, we are able to identify conserved genes with tissue-specific expression patterns. To identify cap-upregulated transcription factor genes, we examined data from Krizsán et al. (2019)^15^. The transcription factor 374546 in *C. cinerea* A43mutB43mut pab1-1#326 (*C. cinerea AmutBmut1*). stood out as a cap-upregulated gene in *C. cinerea*. It was upregulated in the cap in the young fruiting body stage compared to other tissues in the same developmental stage and other stages^15^ (Supplementary Fig. 1). Orthologous genes in three other Agaricomycetes species (*Laccaria bicolor*, *Armillaria ostoyae*, *Lentinus tigrinus*) were also upregulated in caps, relative to other tissue types (Supplementary Fig. 1) based on published data^15–18^.

The gene was deleted in *C. cinerea* by using two CRISPR/Cas9 ribonucleoprotein complexes^19^. Two gRNAs were designed at both ends of the target gene (Supplementary Fig. 2B), which allowed the deletion of a 1,907 bp gene fragment theoretically (Supplementary Fig. 2). The transformants were screened by colony PCR using a primer pair (gcd1_GF/gcd1_GR) designed targeting both ends of the gene (Supplementary Table 1). Transformants that did not show wild-type (WT) bands (Supplementary Fig. 2C) in the colony PCR were further confirmed by RT-PCR using cDNA as the template and gcd1_RT_1F/gcd1_RT_1R as examining primer (for primers see Supplementary Table 1, for the result see Supplementary Fig. 2E). We obtained two independent mutants, which displayed similar phenotypes, so we selected one for the subsequent analyses. To verify that the resulting phenotypes are indeed the consequence of the deletion of the targeted loci, we complemented the *gcd1* deletion by ectopically introducing a FLAG-tagged version of *gcd1* into Δ*gcd1* mutant that resulted the strains cΔ*gcd1* (*‘*c’ denotes complementation). Complementation of the deletion mutant was firstly confirmed by colony PCR (Supplementary Table 1 and Supplementary Fig. 2F). The presence of the GCD1 protein was subsequently verified by western blot analysis of nuclei isolated samples from the FLAG-tagged cΔ*gcd1* strain (Supplementary Fig. 3).

### The deletion of gcd1 led to aberrant fruiting bodies

We analyzed vegetative and sexual phenotypes for the Δ*gcd1* deletion mutants. The morphology of the vegetative mycelium of the Δ*gcd1* strain was similar to that of WT, though its growth rate decreased by 9.09% (two-way ANOVA, *P* = 0.0003) (Supplementary Fig. 4). Reintroduction of the WT *gcd1* gene into the deletion strain nearly fully restored vegetative growth rate (0.88% decrease, two-way ANOVA, *P* = 0.356) (Supplementary Fig. 4A and 4B).

We observed much stronger phenotypic differences during fruiting body development. In synchronized hyphal knot induction^20^, both the Δ*gcd1* and the WT strains were able to form a regular hyphal knot ring after two hours of light induction followed by 24 hours of dark incubation. The WT hyphal knots developed into primordia with well-defined cap, gill, stipe and universal veil (Fig. 1A, C), and eventually to mature fruiting bodies (Fig. 1B). In contrast, the Δ*gcd1* strain produced irregular, spherical to donut-shaped primordia with a smooth surface that apparently lacked universal veil (Fig. 1A, C). Most Δ*gcd1* primordia developed a lower, disc-shaped base and a spherical apical region, which is discernible also in cross-sections (Fig 1A, C). The inner structure of the Δ*gcd1* primordia displayed disturbed differentiation with cap-like and stipe-like structures. The Δ*gcd1* strain also developed slower, its primordia developed from 1 to 5 mm within seven days, whereas the WT strain completed sporulation and deliquescence within this time frame (Fig. 1B). Reintroduction of the flag tagged *gcd1* allele successfully rescued this phenotype (Fig. 1B). We also observed that Δ*gcd1* primordia continued to grow until the agar plate desiccated. This suggests that mechanisms that control size in WT are not fully functional in the Δ*gcd1* strain. This also suggests that the Δ*gcd1* strain does not switch to growth by cell expansion, unlike WT *C. cinerea*, which, after the ∼P2 stage mostly grows by expanding its cell volume. Rather, the Δ*gcd1* strain might maintain active growth and cell proliferation (see also RNA-Seq results below).

**Fig. 1.**
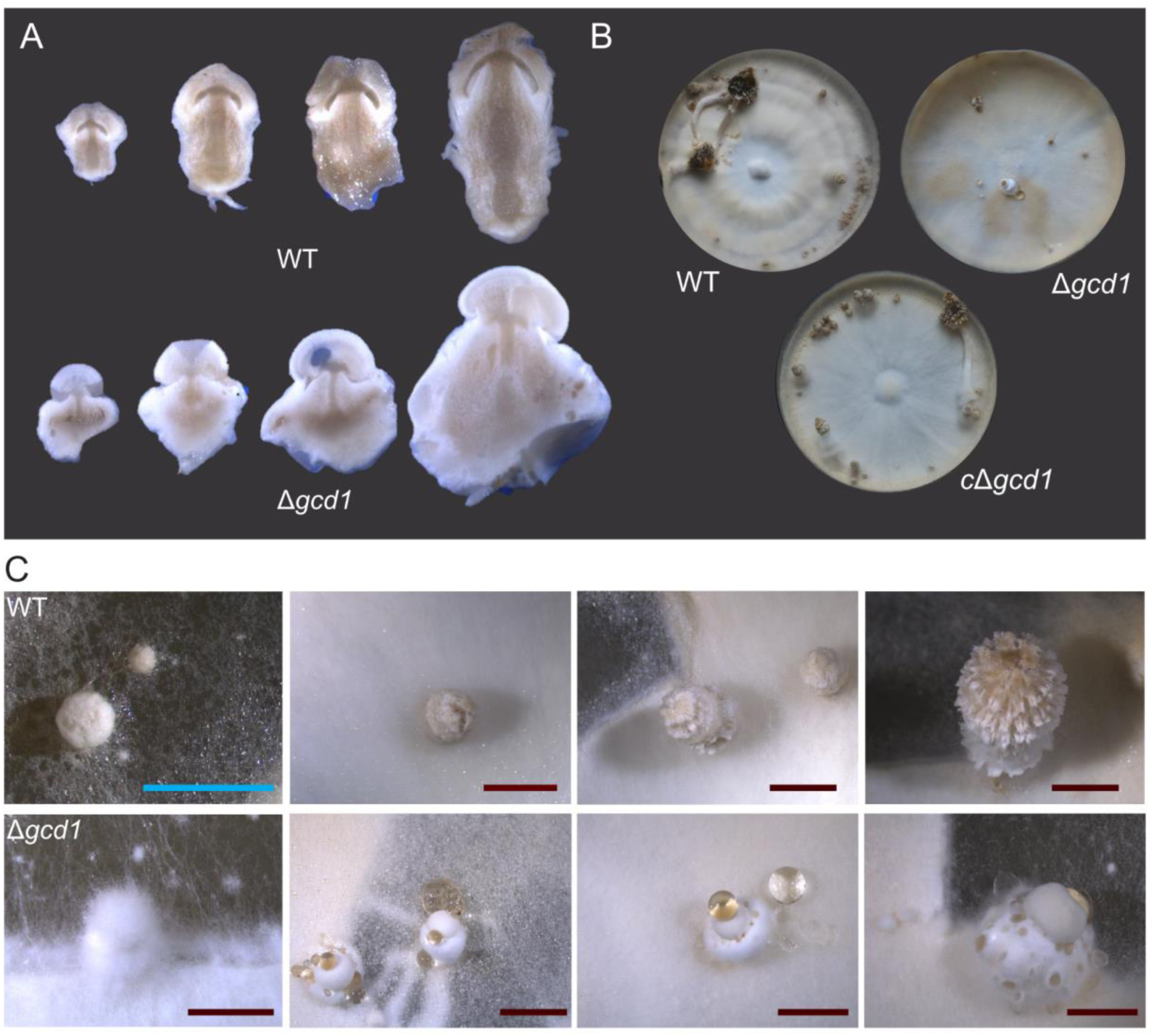
Fruiting bodies of WT and Δ*gcd1 C. cinerea* strains. A, cross-sections of developing fruiting bodies with similar sizes. Strains were grown on YMG medium with halved glucose content at 28°C; B, colonies 7 days post-light induction, with WT and cΔ*gcd1* completing its development whereas Δ*gcd1* strain was in a primordium stage; C, successive developmental stages of WT and Δ*gcd1* strains.

### The Δ*gcd1* mutant lacks veil tissues

Histological sections of primordia with a size of ∼1-5 mm from both WT and Δ*gcd1* mutants revealed problems in primordium development (Fig. 2). The WT strain (Fig. 2A) differentiated several cell types (Fig 2B Panel A-E), with enlarged cells in the external nodulus (E) and universal veil (A). Stipe cells (D) were aligned in parallel, while gill cells (C) were arranged in a palisade between the cap and the stipe. The Δ*gcd1* mutant (Fig. 2A) showed a reduction in cellular differentiation (labeled F-J). We recognize rudiments of cap (G), gill (H), stipe (I) and nodulus (J), however, these are malformed and often topologically misplaced relative to their position in WT primordia. At the cap surface, cortical cells (F) are small and narrow, they resemble cap cells, and there is no evidence of cell expansion as expected for universal veil cells. In older Δ*gcd1* primordia the nodulus enlarges disproportionately, leading to a basal disc-like appearance.

**Fig. 2.**
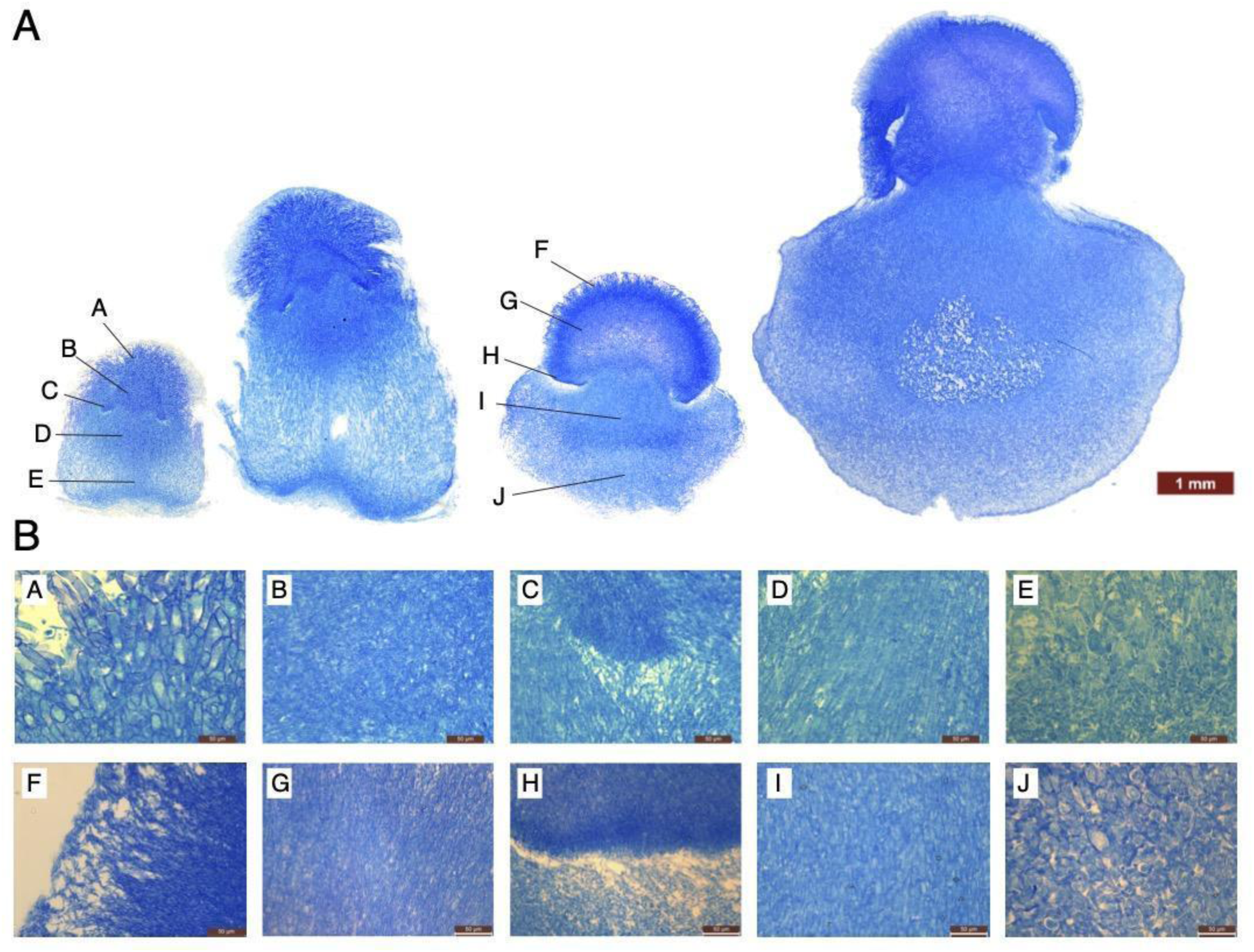
Histology of WT and *gcd1* mutant strains. A, histological sections of primordium stages from WT and Δ*gcd1* stained with methylene blue. B, distinct tissue regions from WT (panel A-E) and Δ*gcd1* (panel F-J). Tissues are labeled as follows: universal veil (A), cap (B), gills (C), stipe (D), nodulus (E); The labeling in Δ*gcd1* (panels F–J) corresponds to the respective regions in WT. Scale bar = 50 µm

To verify tissue identities at the molecular level, we performed quantitative reverse transcription PCR (qRT-PCR) analysis on 3–4.5 mm primordia, using marker genes selected from a high-resolution tissue-specific RNA-Seq dataset (Supplementary Table 2; Supplementary Fig. 5; Supplementary Table 1 for primer sequences). Primordia from both the WT and Δ*gcd1* mutants were dissected into upper cap-like and lower stipe-like regions (Fig. 3). The universal veil marker gene (ID: 470115) showed no detectable expression in either part of the Δ*gcd1* mutant, suggesting the absence of veil in the mutant (Fig. 3). In contrast, the cap marker gene (ID: 422976) exhibited high expression in the cap-like region in WT and reduced to about one-third expression of WT in the Δ*gcd1* mutant (Fig. 3). We observed that the gill marker (ID: 391063) gene has some expression in both the cap-and stipe-like region in both the WT and Δ*gcd1* mutant. Gills occupy a small mass ratio of the fruiting body, this may affect the detection of expression from the bulk-tissue samples collected, nevertheless, the result suggested the presence of gills in the mutant, as also deduced from microscopy observation of sections (Fig. 2B Panel H). The stipe marker gene (ID: 529780) displayed a low and comparable expression level between the WT (Normalized Ct values are between 7.40 to 10.65, Supplementary Table 2) and Δ*gcd1* (Normalized Ct values are between 5.38 to 10.63, Supplementary Table 2) mutants. These results suggested an absence of universal veil cells in the Δ*gcd1* mutant and that cap-, gill- and stipe-like cells are likely present in the Δ*gcd1* mutant. The absence of veil makes Δ*gcd1* mutant primordia gymnocarpic, as opposed to the angiocarpic primordia of the WT. Based on this, we designated *C. cinerea* 374546 as *gcd1*, referring to the gymnocarpic development.

**Fig. 3.**
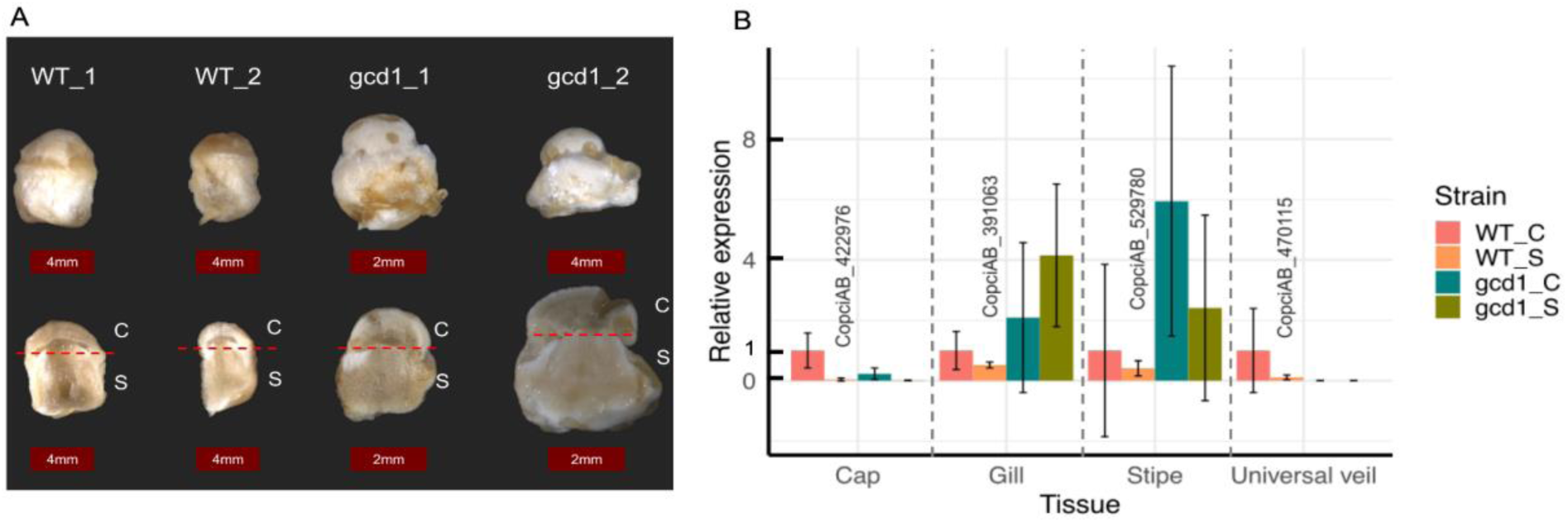
Quantitative RT-PCR analysis of tissue marker genes by using primordia from the WT and Δ*gcd1* mutant. A, primordia used for the qRT-PCR analysis. Primordia were dissected along the lines indicated by red dashed lines into two parts: the upper part (using the letter C to represent), including cap-, gill-, and universal veil-like structures, and the lower part (using the letter S to represent), including stipe-like structures. B, relative expression levels of tissue marker genes in WT and Δ*gcd1* mutant strains, determined by qRT-PCR. Expression is normalized to the WT cap sample. Three technical replicates per RNA sample and two biological replicates were averaged, and the standard deviation is shown on bars.

To the best of our knowledge, morphologies similar to what we observed for the Δ*gcd1* mutant have not been reported in the literature before and only a few comparable mutants are available. The *ich1* mutant in *C. cinerea*^21^, also exhibited developmental problems and eventually an arrest; it produced barrel-shaped fruiting bodies but failed to develop a cap. Földi et al. (2024)^22^ reported the *snb1* mutant in *C. cinerea*, which formed spherical, rudimentary fruiting bodies that failed to differentiate caps, stipes, and gills. Both the *snb1* and *ich1* phenotypes are, however, different from that of the Δ*gcd1* mutant, suggesting that these genes may regulate different aspects of fruiting body development. The *gcd1* mutant seems to develop more tissue types than either previously reported mutant, though in aberrant or rudimentary form, which perhaps preclude continued development towards sporulation.

### GCD1 is a conserved member of the Con7 subfamily of C2H2 zinc finger transcription factors

The *gcd1* gene encodes a 670 amino acid polypeptide which contains one C2H2-type zinc finger domain (IPR013087). Based on its domain content it belongs to a Con7 subfamily (IPR039327) which was first identified in *Pyricularia oryzae*^23^ (as *Magnaporthe grisea*). We analyzed the conservation and gene copy number distribution of *gcd1* by using two datasets, one comprising 655 published genomes (G655) from MycoCosm^24^ and NCBI (Fig. 4), (Dikarya and non-Dikarya fungi, with Nucleariida and Holozoa as outgroups) and a dataset of 189 species with higher resolution in the Agaricomycetes^8^.

**Fig. 4.**
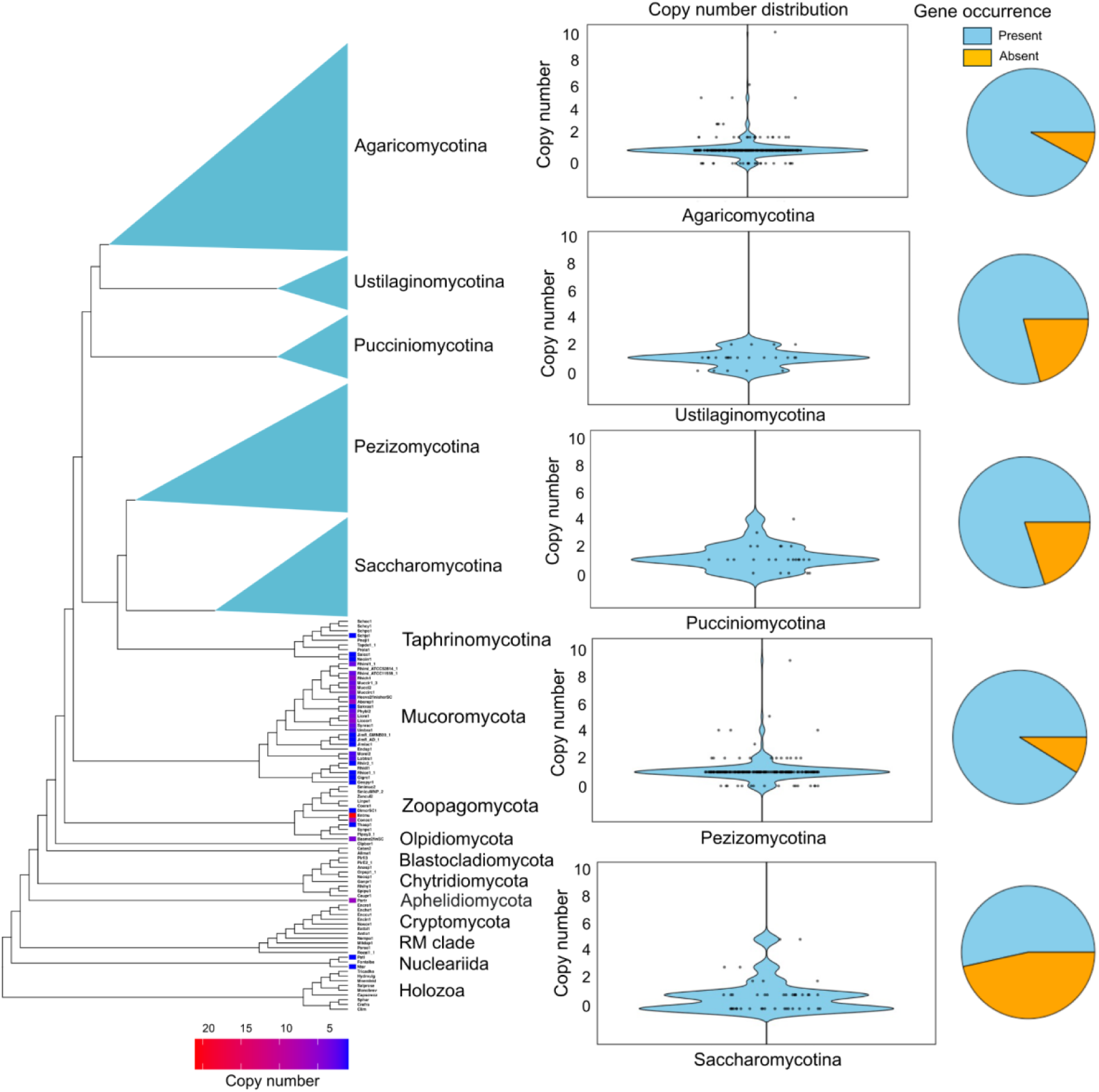
Phylogenetic distribution of *gcd1* and its copy numbers in G655 dataset. For Taphrinomycotina species and each non-Dikarya species, the copy number of the *gcd1* gene is shown next to the corresponding leaves. Violin plots display the distribution of *gcd1* gene copy numbers within each major clade illustrating variation and central tendency. Pie charts summarize the proportion of species within each clade that possess the *gcd1* gene.

Our analysis indicates that the Con7 subfamily of transcription factors (i.e. GCD1 homologs) are mostly encoded by a single gene in species that possess it. This was true both in the 655-species (Fig. 4.) and the 189-species datasets (Supplementary Fig. 6). We found the largest variance in copy numbers in the Mucoromycota, where certain species have >10 copies of the encoding gene.

We found that Con7/GCD1 homologs are prevalent in Agaricomycotina (92%), Pezizomycotina (91%), Ustilaginomycotina (79%) and Pucciniomycotina (80%). In contrast, only 53% of the examined Saccharomycotina species and 33% of the examined Taphrinomycotina species possess this subfamily. Notably, the fruiting-body-forming *Neolecta irregularis* (within Taphrinomycotina), despite being nested among predominantly yeast-like taxa, possesses a GCD1 homolog. GCD1 homologs were rarely present in the examined Zoopagomycota species (17%) and were not detected in the Holozoa clade and the Rozellida-Microsporidia clade, whereas nearly all Mucoromycota (88%) have GCD1 homologs. The C2H2 TFs have undergone significant expansion during fungal evolution^25^. Con7/GCD1, as a member of this family, likely arose during this diversification. Its distribution, in fungi and absence in non-fungal lineages, indicates that Con7/GCD1 homologs form a fungal-specific subfamily of C2H2, reflecting fungal-specific evolutionary innovation in transcriptional regulation. To assess its conservation within Agaricomycotina, we performed a phylogenetic analysis by using GCD1 homologs identified in 189 published genomes^8^. The gene tree revealed that GCD1 is present in a single copy in most species (Supplementary Fig. 6), in agreement with the comparative genome dataset above. The gene tree approximately resolved main clades in the Agaricomycotina, however, it included several unstable nodes and generally low bootstrap support values. This is due to the high variability of the amino acid sequences in the multiple alignment, which could not be reliably resolved despite multiple alignment strategies tested. Nevertheless, our analysis indicates that the GCD1 is broadly found across the fungal kingdom, is very likely fungal-specific and is encoded by a single copy gene in most species.

### Transcriptomic analysis of the Δ*gcd1* and WT strains

The above results indicate that the Con7 subfamily occurs much more broadly (across the Dikarya) than gymno- and angiocarpic fruiting bodies do, suggesting that it regulates more basic functions than fruiting body type. It follows that gymnocarpy in the mutant is then an indirect consequence of the deletion of *gcd1*. To understand how the deletion causes the observed phenotypic differences, we performed transcriptomic analysis by using the hyphal knot stage. We opted for this stage because in the secondary hyphal knot there is minimal morphological divergence between WT and Δ*gcd1* strains, which allows us to detect direct effects of *gcd1* deletion better than samples which differ morphologically. Based on previous transcriptomic data^15^, *gcd1* has a very low expression level in the vegetative mycelium but is highly upregulated in secondary hyphal knots (Supplementary Fig. 1A) which suggests its regulatory impact can be studied at these early stages.

We performed RNA-Seq on the hyphal knot ring from WT and Δ*gcd1* strains after 2 hours of light induction followed by 24 hours in darkness (Fig. 5A). At this stage both WT and Δ*gcd1* have hyphal knots with some internal differentiation^9^, and at this stage we believe there is no major structural difference between the two sample types that would introduce unnecessarily high numbers of unspecific gene expression differences into the analyses. Pearson correlation coefficients showed good consistency for both WT and Δ*gcd1* biological replicates (Fig. 5B). The transcriptomic analysis revealed 935 differentially expressed genes (DEGs) (Supplementary Table 3a and 3b) with a fold change greater than 1 and a BH-adjusted p-value < 0.01 (Wald test). We applied a low fold-change threshold to identify DEGs due to the presence of vegetative mycelium in the samples, which may reduce the relative abundance of transcripts associated with hyphal knots. Of the 935 DEGs identified, 350 were upregulated, while 585 were downregulated (Fig. 5C, Supplementary Table 3a and 3b).

**Fig. 5.**
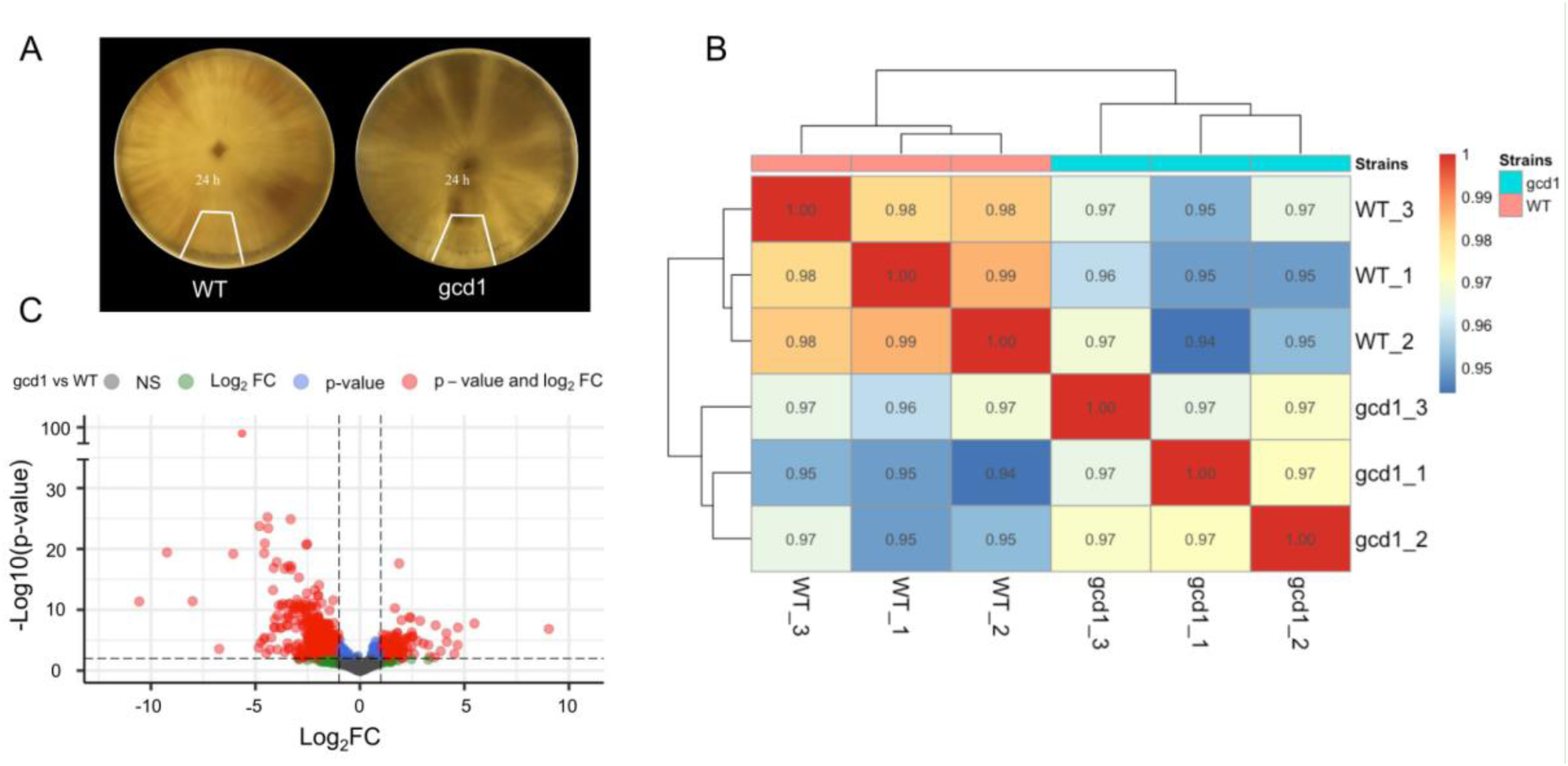
RNA-Seq analysis of the hyphal knot stage of the *Δgcd1* strain versus WT. A, sampling scheme for RNA-Seq. The sampled region is delineated by a white line; B, heatmap of sample correlations. The numbers in the cell are the Pearson correlations coefficients across samples; C, Volcano plot of the differentially expressed genes. The gray and green color represents non-significant differentially expressed genes (DEGs) with an FDR adjusted p-value >= 0.01 (Wald test). The blue color indicates genes with an FDR-adjusted p-value < 0.01 (Wald test) and a log2 fold change (log2FC) between 0 and 1. Red denotes significant DEGs with an FDR-adjusted p-value < 0.01 (Wald test) and a log2FC greater than |1|.

DEGs included few experimentally characterized genes, which drove us to functionally characterize them using computational tools. Exceptions include the cospin *pic1*^26^ (367957) and *keps*^27^ (426342 and 503649) among downregulated genes, and *mep3* fungalysin^28^ (26022), *cfs1*^29^ (447925) and the hydrophobin *CoH2*^30^ (468111) among upregulated genes. To computationally characterize DEGs, we used Gene Ontology terms and tested the overlap of Δ*gcd1* DEGs with five published gene lists^15,31^ describing fruiting body initiation, mycelium type, starvation and light response in *C. cinerea* (Supplementary Table 6). Hereafter we discuss results of these analyses, with a focus on downregulated genes, because we expect these to comprise morphogenesis-related genes directly or indirectly affected by GCD1. We also focus on upregulated genes, which give some clues on potential mechanisms explaining the growth of Δ*gcd1* primordia.

Results of Gene Ontology (GO) enrichment analyses are summarized in Fig. 6 and Supplementary Table 4. Overall, 32.14% and 60.57% of down- and upregulated DEGs are associated with any GO term, respectively, compared to 47.17% of all protein coding genes in the genome. This represents a significant depletion of GO terms in downregulated DEGs (Fisher’s Test p-value = 5.67 × 10⁻^14^) (Supplementary Table 4a) and suggests that the functions of downregulated DEGs in the Δ*gcd1* mutant are particularly poorly annotated functionally compared to the genome-wide average in *C. cinerea*. In contrast, a higher ratio of upregulated DEGs had GO annotations (Ratio: 60.57%; Fisher’s Test p-value 3.88 × 10^−7^) than all genes in the genome (Supplementary Table 4a), suggesting that upregulated DEGs include better annotated biological functions, and perhaps more conserved genes.

**Fig. 6.**
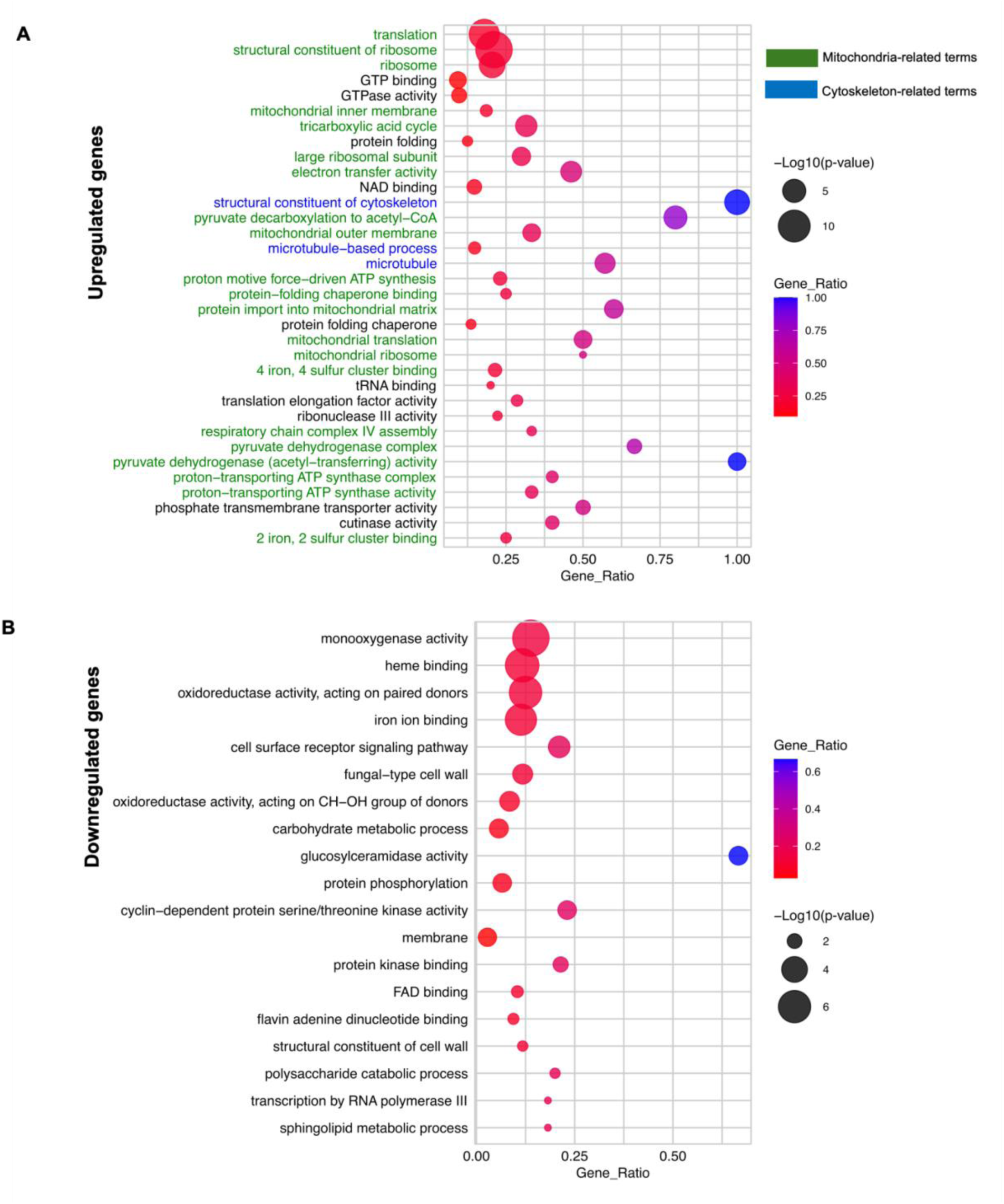
GO enrichment analysis of the Upregulated DEGs (A) and Downregulated DEGs (B).

### Downregulated genes

Among the downregulated genes, a total of 19 GO terms were significantly enriched (Fisher’s exact test, p-value ≤ 0.05), with 92 genes annotated to these enriched GO terms. Of these we here highlight those related to the fungal cell wall (FCW) (carbohydrate metabolic process, polysaccharide catabolic process, glucosylceramidase activity, fungal-type cell wall and structural constituent of cell wall) and cell signaling pathways (protein phosphorylation, protein kinase binding) (Supplementary Table 4b).

In association with the cell wall we identified two putative chitin deacetylase genes in the CE4 family (440521, FC_wt/Δ*gcd1*_: 4.05; 518489, FC_wt/Δ*gcd1*_: 2.53), a GH18 chitinase (488885, FC_wt/Δ*gcd1*_: 2.95), four glucanase-encoding genes (366990, FC_wt/Δ*gcd1*_: 3.01; 376074, FC_wt/Δ*gcd1*_: 2.40; 459642, FC_wt/Δ*gcd1*_: 1.80; 524741, FC_wt/Δ*gcd1*_: 1.83) belonging to GH16 and one belonging to GH12 (361787, FC_wt/Δ*gcd1*_: 2.65), which are likely involved in glucan remodeling. In the context of the complete developmental process of *C. cinerea*, one of the genes (459642) was almost exclusively expressed in the stipe of the mature fruiting body^15^ (Supplementary Table 3b) indicating its potential role in stipe development. Another glucan-related gene (361787) had the highest expression level in caps and gills of the mature fruiting body in *C. cinerea*^15^ (Supplementary Table 3b). Even though we detected these downregulations in the hyphal knot stage, we believe that some of this can explain the morphological defects that we see in the later stages.

We also identified two downregulated genes encoding putatively glucan-related GH30_3 family proteins (372384, FC_wt/Δ*gcd1*_: 2.43; 381637, FC_wt/Δ*gcd1*_: 1.69). One of these (381637) has been functionally characterized in *C. cinerea* (as GH30A) and shown to have endo-1,6-glucanase activity^32^. Its high expression level in mature caps^32^ (see also Supplementary Table 3b) suggested a role in autolysis. In *Lentinula edodes*, its ortholog, LePus30A exhibited endo-β-1,6-glucanase activity against fruiting body cell walls and was suggested to be responsible for senescence^33^. The GH30A gene of *C. cinerea* has also been proposed to play a role in cell wall remodeling by replacing β-1,6-glucan to β-1,3-glucan crosslinking in the stipe base, which may reduce wall extensibility, leading to the cessation of growth^32^. Our results on the Δ*gcd1* mutant indicate potential links between the GH30_3 family and early developmental events in *C. cinerea*.

Given the apparent importance of GH30_3 in fruiting body development, we conducted a phylogenetic analysis of GH30_3 protein sequences and inferred their ancestral copy numbers across using 109 species (G109) including Basidiomycota (104 species) and Ascomycota (5 species) using a previously published dataset^34^. The examined Ascomycota had very few GH30_3 genes (mean: 0.2, 5 species). In the Basidiomycota, GH30_3 genes occur patchily in non-Agaricomycetes clades (mean: 0.9, 15 species), whereas nearly all examined Agaricomycetes had GH30_3 genes (mean: 2.1, 89 species) (Fig. 7). Ancestral copy number inferences revealed several duplications in Agaricomycetes (see arrows in Fig. 7). Consistent with this, early-diverging orders (Cantharellales to Russulales) have on average 0.9 GH30_3 genes, while Polyporales, Boletales and Agaricales possess 2.1, 2.1 and 2.2, respectively. At the scale of the Agaricomycetes these observations indicate higher GH30_3 copy numbers in orders that possess complex fruiting bodies, although at finer scales the correlation with complexity breaks down (e.g. only two genes in *C. cinerea*, but nine in *Fibulorhizoctonia psychrophila).* Nevertheless, the GH30_3 family seems to be expanded in Agaricomycetes. In addition, one of the genes from GH30_3 family (381637) has been reported to be developmentally regulated among 11 examined species with complex multicellular fruiting bodies^10^. Taken together, we speculate the expansion of genes from GH30_3 may be linked to the evolution of fruiting bodies.

**Fig. 7.**
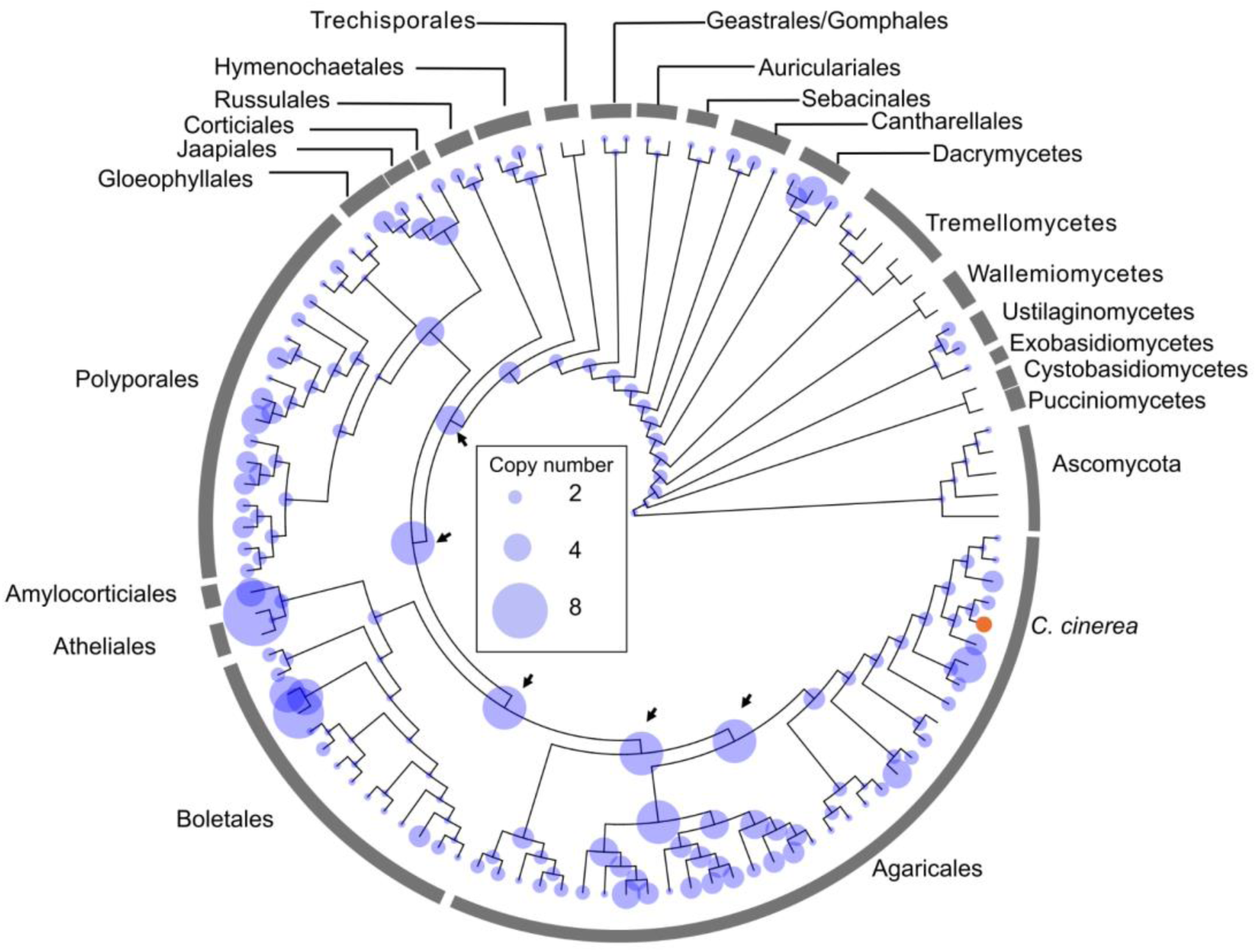
Species tree of G109 dataset showing extant and inferred ancestral copy numbers of the GH30_3 family in 109 species. *C. cinerea* is highlighted with orange, which has two paralogs (381637 and 372384). The phylogenetic tree and gene family evolution analysis was obtained from a published dataset^34^.

Another group of four downregulated genes linked to the FCW encode hydrophobins (361161, FC_wt/Δ*gcd1*_: 3.85; 413238, FC_wt/Δ*gcd1*_: 5.61; 419467, FC_wt/Δ*gcd1*_: 1.50; 427537, FC_wt/Δ*gcd1*_: 11.05). Hydrophobins have been extensively studied in fruiting body development in mushroom-forming fungi^35–38^, as well as in pathogenicity in plant-pathogenic fungi^39^. In plant-pathogenic fungi, deletion of *gcd1* also led to a downregulation of hydrophobin genes^39^.

The next significant functional enrichment was associated with signaling pathways (protein phosphorylation (GO:0006468) and protein kinase binding (GO:0019901)) (Fig. 6B). In these categories we found 20 downregulated kinases, of which 13 belong to the fungal specific FunK1 kinases and two to Tyrosine kinase-like (TKL) kinases (Supplementary Table 5). Both FunK1 and TKL kinases have expanded significantly in Agaricomycetes^15^. Eleven out of 20 downregulated kinase-encoding genes were developmentally regulated in Agaricomycetes fruiting bodies^15^, indicating a dysregulation of regulatory signaling pathways may be associated with defective fruiting body development in our mutant.

Gene set enrichment analyses indicated that downregulated DEGs showed significant overlap with genes related to fruiting body initiation, cap-upregulated genes, aerial-mycelium specific genes as well as gene sets defined as late light response and core starvation^15,31^ (Fig. 8B). Of these, a high overlap with cap-upregulated genes (gene number/DEGs: 124/585, FDR adjusted p-value = 1.95×10^−13^, Hypergeometric test; Supplementary Table 6) is of particular interest, given the malformation of caps and the lack of veil in the mutant. We identified two downregulated genes related to defense; the fungal peptidase inhibitor *pic1*^26^ (367957, FC_wt/Δ*gcd1*_: 27.99, MEROPS family I66) and a thaumatin-encoding gene identified as defense-related protein by Plaza et al. (2014)^40^ (496344, FC_wt/Δ*gcd1*_: 2.74). *Pic1* is a cospin that functions as an effector of a fungal defense mechanism against fungivorous insects, whereas thaumatins are involved in antimicrobial defense against bacteria and fungi^26^. Given the defense-related function of the above two genes, we mined ultra-low input RNA-Seq data (Supplementary Table 6k) and found that these two genes exhibited very high in the universal veil (Supplementary Fig. 7), suggesting a role at the veil surface. The downregulation of this gene aligns with the absence of a universal veil in the Δ*gcd1* mutant.

**Fig. 8.**
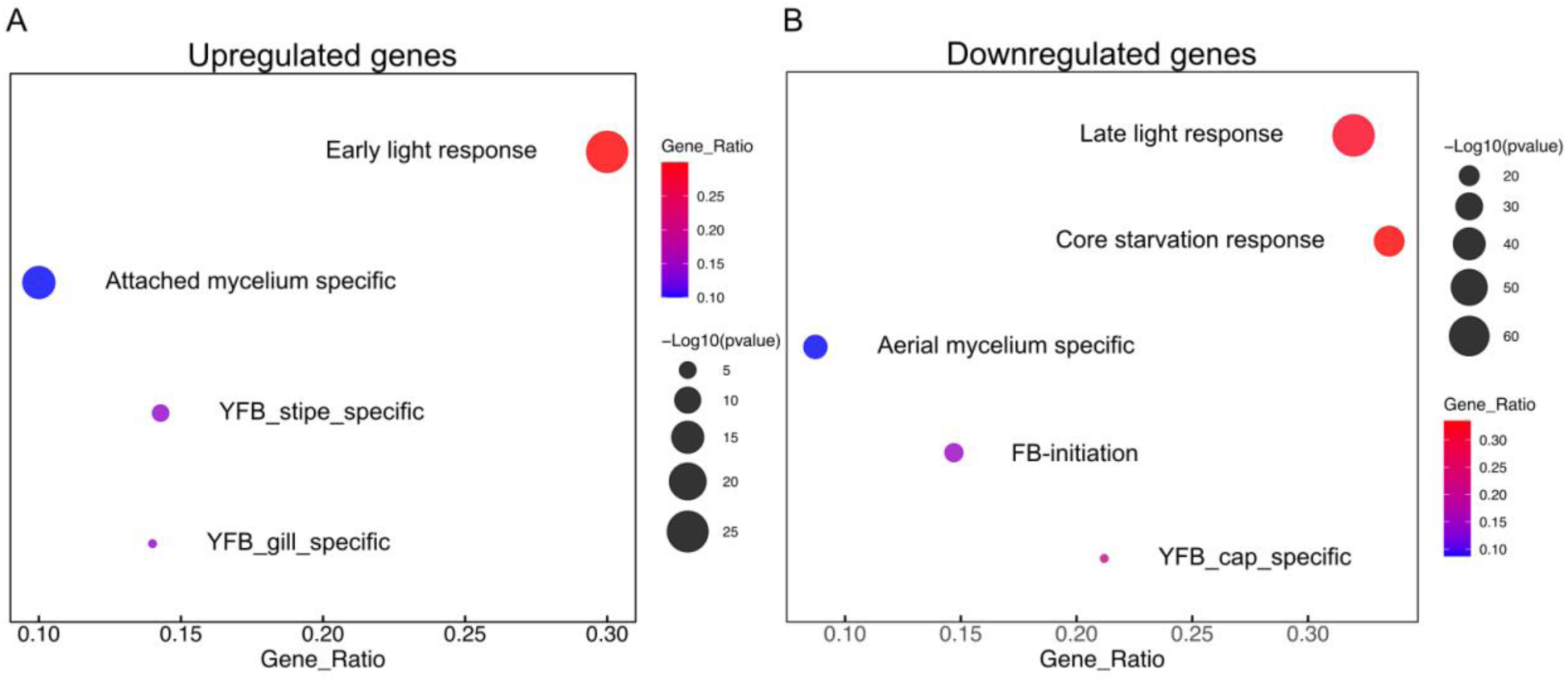
The custom gene set enrichment analysis of upregulated DEGs (A) and downregulated DEGs (B) with a significance threshold of FDR adjusted p-value (Wald test) < 0.01.

We also found glucose receptor-encoding genes (*Gpr1 in S. cerevisiae*^41^), that overlap between *gcd1* downregulated DEGs and the cap-specific gene set. Out of five *Gpr1* genes predicted in *C. cinerea*, three are downregulated (426382, FC_wt/Δ*gcd1*_: 3.57; 559335, FC_wt/Δ*gcd1*_: 2.13; 263790, FC_wt/Δ*gcd1*_: 2.17) and one was postulated to be a cap-upregulated gene^15^. Gpr1 is a G protein-coupled receptor (GPCR) that serves as a primary sensor for extracellular glucose in fungi^42,43^, although there is evidence that it can respond to other molecules too^44^. The broad downregulation of glucose receptors in the mutant may signify changes in sensing nutrient availability or other signaling mechanisms.

FB initiation genes were identified by Krizsán et al. (2019)^15^ as those whose expression increased at least fourfold at the hyphal knot and stage 1 primordia, relative to vegetative mycelium. Among 585 downregulated DEGs, there are 86 genes (15%) overlapping with this category (FDR adjusted p-value = 2.91×10^−18^, Hypergeometric test; Supplementary Table 6), while only 19 upregulated DEGs (5%) could be classified to this group (FDR adjusted p-value > 0.05, Hypergeometric test). This underscores that the genes downregulated in Δ*gcd1* are associated with early developmental processes leading to the emergence of tissue types.

Beyond genes identified by GO and gene set enrichment analyses, we found a notable downregulation of three Pcl1/2-like cyclin homologs (1001791, FC_wt/Δ*gcd1*_: 2.93; 1001807, FC_wt/Δ*gcd1*_: 2.93; 384611, FC_wt/Δ*gcd1*_: 2.71) Cyclins are conserved and generally not expressed dynamically during fruiting body formation^15^. Pcl1/2-like cyclins partner with the cyclin-dependent kinase Pho85 to regulate the cell cycle, polarity and morphogenesis in *S. cerevisiae*^45^, and affect germination, cell division, and sporulation in filamentous fungi^46,47^. The role of Pcl1/2 cyclins in fruiting body development is unknown, nevertheless, we hypothesize they participate by regulating the cell cycle.

### Upregulated genes

Upregulated genes are associated with 34 significantly enriched GO terms (Fisher’s exact test, p-value ≤ 0.05). Given that many terms were clearly mitochondrion-related, we examined the genes within each term and categorized them as mitochondria-related or other, using subcellular localization predictions for classification (Fig. 6; green vs. other colors). Among these mitochondrion-related GO terms, we found ones referring to organelle parts, mitochondrial ribosomes, and respiration/metabolism, which overall suggest increased mitochondrial activity in the mutant. In other mitochondria-related GO terms, we identified proteins involved in energy production (GO:0006086 “pyruvate decarboxylation to acetyl-CoA”; GO:0006099 “tricarboxylic acid cycle”; GO:0015986 “proton motive force-driven ATP synthesis”).

The GO analysis highlighted several ribosomal terms. To check if these correspond to mitochondrial or cytoplasmic ribosomes, we annotated the entire ribosomal apparatus in *C. cinerea* (Supplementary Table 7a). We found 92 ribosomal genes, 67 mitochondrial ribosomal genes and 42 ribosome biogenesis genes. In the context of the Δ*gcd1* upregulated genes, we found a significant overrepresentation of mitochondrial ribosomal genes (DEGs/All annotated, 34/67, Fisher’s exact test p-value 7.0 × 10^−36^) (Supplementary Table 7b). We also observed three GO terms related to the cytoskeleton. They correspond to four tubulin genes, with two beta-tubulins (393528, FC_Δ*gcd1*/wt_: 1.59 and 519006, FC_Δ*gcd1*/wt_: 2.18) and two alpha-tubulins (176636, FC_Δ*gcd1*/wt_: 1.69 and 359665, FC_Δ*gcd1*/wt_: 1.74). Tubulins form the core structural components of microtubules, which are known to mediate organelle positioning and intracellular transport, including mitochondrial dynamics^48^. We hypothesise that tubulin upregulation also corresponds to a higher number of mitochondria and thus increased need for positioning.

Taken together, we interpret the GO results of upregulated genes as a broad signal for increased expression of mitochondria-related genes, which in fungi may reflect increased energy demands and stress responses^49–51^. In our case, we hypothesise that this may suggest elevated energy consumption during the hyphal knot stage of the mutant, which may be caused by inadvertent misregulation as a result of the gene knockout in the mutant.

## Conclusions

In this study we described a novel transcription factor, GCD1 that is required for fruiting body morphogenesis in the model mushroom *Coprinopsis cinerea.* We show that its deletion leads to disturbed morphogenesis and the lack of veil tissues, resulting in a gymnocarpic (open) fruiting body. We showed that GCD1 belongs to the fungal-specific Con7 subfamily of C2H2 zinc-finger transcription factors, which is conserved and harbors a single gene in most Dikarya species, suggesting high functional conservation.

Members of the Con7 subfamily have been primarily investigated in plant pathogenic fungi. These genes have been functionally characterized in *Pyricularia oryzae*^23^, *Fusarium oxysporum*^39^, *Parastagonospora nodorum*^52^, *Fusarium graminearum*^53^ and two *Colletotrichum* species^54^ to date, plant pathogens in which the deletion of the gene yielded reductions in pathogenicity and virulence as well as diverse morphogenetic (e.g. conidiation) defects. A consensus across these studies is that Con7 homologs regulate pathogenicity and morphogenesis; with mechanistic insights suggesting that the defects in the former indirectly result from defects in morphogenesis. In terms of morphogenesis, Con7 homologs have been linked to conidiation, the development of sexual and infection structures (e.g. appressoria), but not vegetative growth^53^. The latter agrees with our observations in *C. cinerea*, where we found that *gcd1* is hardly expressed in vegetative mycelium and its deletion led to a marginal difference in vegetative growth rate. On the other hand, it is highly expressed in sexual fruiting bodies and we documented a significant impact of its deletion on the latter. Based on these observations we speculate that the regulation of non-vegetative morphogenesis may be a general property of the Con7 subfamily of fungal transcription factors.

We might ask, then, whether the similar role of Con7 homologs across species manifest in shared target genes. Δ*gcd1* transcriptomic results and target genes of Con7 homologs in other species show some distinct similarities. Several studies reported that cell wall synthesis and remodeling enzyme genes (e.g. chitinases) are Con7-dependent^23,39,52,53^. Our RNA-Seq data indicated a downregulation of CE4 chitin deacetylases, GH18 chitinases and multiple glucanases in *C. cinerea,* raising the possibility that these are targets of GCD1, although direct evidence is missing in our case. These similarities suggest that, like Con7 homologs in ascomycetes, GCD1 is a primarily morphogenetic transcription factor in mushroom-forming fungi.

Transcriptome analyses provided insights into the altered gene expression that characterizes the Δ*gcd1* phenotype. Downregulated genes included ones related to the fungal cell wall and encoding various chitinases and glucanases, several hydrophobins, kinases (FunK1 and tyrosine kinase-like kinases), cyclins among others. We demonstrated that the GH30_3 family, which encodes glucanases, has undergone an evolutionary expansion in mushroom-forming fungi, suggesting a link to fruiting body development. Upregulated genes, on the other hand, were predominantly related to energy metabolism. We speculate this is linked to the unregulated growth of the mutant, which eventually results in considerably larger primordia than in the wild type. However, as these patterns were detected at the hyphal knot stage, which did not show any morphological differences with wild type strains, the upregulation of mitochondrial genes may be a potentially direct consequence of the deletion of *gcd1*.

GCD1 is currently the single functionally studied member of the Con7 subfamily in the Agaricomycetes. The closest homolog of GCD1 with functional information is ZFC3 in *Filobasidiella (Cryptococcus) neoformans*, however, this has only been examined in the context of pathogenicity^55^. Future studies should investigate more species to more thoroughly understand its role in fruiting body morphogenesis and beyond. Speculations of similarly important morphogenetic roles in other species are underpinned by the expression dynamics GCD1 homologs of other species comparable to that in *C. cinerea*. Another avenue for future research is investigating the regulatory network of GCD1, both upstream, its regulators, and downstream, its target genes. As mushroom functional genomics improves, these will come more within reach, even for proteins like GCD1 which appears hard to handle using functional genomics essays.

Taken together, this study demonstrated that GCD1 is required for proper fruiting body morphogenesis in a model basidiomycete. The mutant phenotype, phylogenetic conservation of the protein sequence and the transcriptomic results combined with data on other Con7 homologs suggest that this subfamily of transcription factors is broadly related to morphogenesis in the fungal kingdom. While this assertion will need to be tested in the future, the sequence conservation and function of the Con7 subfamily seems to transcend the >500 million years of evolution since the Asco- and Basidiomycota diverged from each other.

## Acknowledgements

The authors acknowledge support by the National Research Development and Innovation Office (Grant No. OTKA 142188), the European Research Council (grant no. 101086900 to L.G.N.).

## Author contributions

H.W. and L.G.N. conceived the study. H.W., Z.M., M.V., X-B.L., B.H., Z.H., E.A., A.F., Z.K., and Z.L., performed the wet-lab experiments. H.W., Z.M., B.H. and L.G.N processed the bioinformatics data. H.W., Z.M., and L.G.N analyzed the data and prepared figures. H.W., Z.M., and L.G.N wrote the original manuscript. All authors have read and approved the manuscript.

## Materials and Methods

### Strains and cultures

The homokaryotic strain of *C. cinerea* A43mutBmut pab1-1# 326 strain was cultured in the yeast extract- and malt extract-glucose (YMG) medium at 37°C for 6-7 days under constant light conditions to generate oidia. For fruiting body induction, we followed a synchronized process as described previously^20^. Strains were cultured on the YMG agar medium with half the glucose content (0.2% wt/vol) and incubated under constant darkness at 28°C for 5-6 days until the mycelium reached ∼1 cm from the edge of the Petri dishes. The Petri dishes were then transferred to white light under 28°C for 2 h, after which they were placed back to the dark for 24 h, followed by 12-h light/12-h dark period at 28°C.

To determine the mycelial growth rate on agar plates, small agar plugs (d = 5 mm) of the WT, Δ*gcd1*, and cΔ*gcd1* strains were inoculated onto YMG (0.2% glucose) medium and incubated at 37°C in the dark, with 8 replicates per group. Colony diameters were measured every 24 hours up to 96 hours, and the growth rate was calculated as colony growth per hour. Statistical analysis of the differences among these strains was performed using two-way ANOVA.

### Targeted deletion and complementation

Plasmid used for targeted gene deletion includes the pUC19 backbone linearized by primers, upstream and downstream homology arms (1000 bp), and one selection marker para-aminobenzoic acid (PABA) fragment amplifying from the pMA412 vector^56^ Primers used for the amplification of each fragment are listed in Supplementary Table 1, the plasmid map for gene deletion is shown in Supplementary Fig. 2. Plasmid used for targeted gene complementation includes the pUC19 backbone linearized by primers listed in Supplementary Table 1, the target gene promoter region (1000 bp upstream of the translation initiation codon (ATG)), gene coding region and 1000 bp terminator region amplified from the genome of *C. cinerea* and the selection marker of hygromycin B phosphotransferase gene (hyg) was synthesized by using primers listed in Supplementary Table1, promoter and terminator gene are amplified from *C. cinerea* tubulin-encoding gene. To examine the presence of the GCD1 protein, we tagged it with a 3×Flag tag. Primers used for gene complementation are listed in Supplementary Table 1, the plasmid map for gene complementation is shown in Supplementary Fig. 2.

The CRISPR/Cas9 system was used for gene deletion in *C. cinerea*. For the transformation, the ribonucleoprotein (RNP) complex consisting of Cas9 protein and single guide RNAs (sgRNA) designed at both ends of the target gene, together with the pUC19 (PABA)-plasmid constructed above are delivered to the protoplast of *C. cinerea*. The protoplast transformation of *C. cinerea* was performed following the protocol described by Dörnte et al. (2012)^57^ and Pareek et al. (2022)^19^. For the complementation transformation, the pUC19 (hyg)-plasmid was delivered to the protoplasts of *C. cinerea*. The protoplast transformation of *C. cinerea* was similar to that used in gene deletion, with slight modification. Briefly, after delivering the plasmid to the protoplast, the hygromycin B (Thermo Scientific) was added to the petri dishes after 24 hours with the final concentration of 100 µg/mL.

The Δ*gcd1* mutants were screened by genomic polymerase chain reaction (PCR) with the primers listed in Supplementary Table 1. The PCR products amplified by the primer pair gcd1_GF/GR were shown in Supplementary Fig. 2. The size of the PCR product amplified from WT was 1808 bp; however, samples lacking this band were considered mutants, as the gene fragment might have been deleted or multiple plasmid fragment insertions could have resulted in a band too large to be amplified. *gcd1* complementation transformants were screened with the insertion of the target gene fragments starting from the promoter till the end of the terminator. The PCR primers were designed to amplify 3069 bp genomic fragments of *gcd1* complementation transformants (Supplementary Fig. 2).

### RNA-Seq and data analysis

We sampled the hyphal knot ring along with mycelium from the WT strain and Δ*gcd1* stain after 2 hours of light induction followed by 24 hours in darkness. Specifically, the strains were cultured on the same medium used for fruiting, with each Petri dish covered with cellophane. After 5-6 days, Petri dishes were exposed to light for 2 hours and then placed back for another 24 hours in darkness, the samples were collected as shown in Fig. 5A. RNA extraction was performed using the Quick-RNA Miniprep Kit (Zymo Research, USA), cDNA library and pair-end (150 bp) sequencing were conducted by Novogene Co., Ltd (Cambridge, United Kingdom). The raw reads quality was checked by FastQC^58^ (version: 0.12.0) and MultiQC^59^ (version: 1.19). Adapters, low quality bases and short reads were trimmed by bbduk.sh from BBtools 38.92^60^ following a parameter of qtrim=rl, trimq=25, minlength = 40. The trimmed reads were mapped to the reference genome using STAR version 2.6.1a_08-27^61^ The aligned reads were counted for each gene in the genome using the summarizeOverlaps function from the GenomicAlignments v.1.22.1 package^62^ using the “Union” mode. The mapping reads statistics is shown in Supplementary Table 8. Sample clustering by Pearson correlation coefficients was performed with the transformed count data by Variance Stabilizing Transformation (VST) in the DESeq2 package^63^. Volcano plots were generated using the EnhancedVolcano package 1.27.0^64^. Genes with >5 raw counts in ≥2 replicates per sample group were kept for further analysis. Differentially expressed genes (DEGs) were identified by using the R package DESeq2 1.48.1^63^. Genes with Log2FC > 0, BH adjusted p-value < 0.01 (Wald test) were considered as upregulated DEGs, and Log2FC < 0, BH adjusted p < 0.01 (Wald test) were considered as downregulated DEGs.

InterPro (IPR) and GO functional annotations were defined using InterProScan version 5.61-93^65^. The GO enrichment analysis was performed by R package topGO 2.60.1^66^. The weight01 algorithm was used with Fisher statistics, and the significance level was set to ≤ 0.05. Putative CAZymes were identified using dbCAN^67^ and further categorized based on their substrate specificity as described in previous publications^68^. Based on the substrate specificity information, the groups of plant cell wall degrading CAZymes (PCWDEs) and fungal cell wall degrading CAZymes (FCWDEs) were defined by Hegedüs et al. (2025)^31^(available at mushroomdb.brc.hu). The ribosomes and mitochondrial ribosomes were identified based on InterPro (IPR) annotations. Subcellular localization predictions were performed using DeepLoc 2.0^69^.

A custom dataset which contains the gene groups for certain processes was created from previous published data (Supplementary Table 6), including genes related to light response, starvation response, aerial mycelium and attached mycelium and fruiting body initiation and tissue_specific gene^15,31^. The enrichment analysis of the custom dataset was conducted by R package clusterProfiler with the minimal size of genes annotated for testing (minGSSize) was set to 2 and the FDR-adjusted p-value cut-off was set to 0.01^70^.

### Quantitative real-time PCR (qRT-PCR) analysis

RNA extraction was performed using the Quick-RNA Miniprep Kit (Zymo Research, USA). The first-strand cDNA was synthesized with RevertAid First Strand cDNA Synthesis Kit (Thermo Fisher Scientific, USA). The qRT-PCR was performed with PowerUp™ SYBR™ Green Master Mix for qPCR (Applied Biosystems, Weiterstadt, Germany). The amplification program started with an initial UDG activation at 50°C for 2 min, polymerase activation at 95°C for 2 min, followed by 40 cycles of 15 seconds denaturation at 95°C and 1 min annealing at 60°C. The *β*-tubulin gene was used as the internal control. Primers used for the qRT-PCR analysis were listed in the Supplementary Table 1.

### The generation of the expression image heatmap

The expression image heatmap was generated using an in/house script applied to an unpublished tissue-specific RNA-seq dataset (Supplementary Table 2 and Supplementary Table 6k). Log2-transformed normalized counts were used for the heatmap, and the color scale was normalized from 0 to 1 to represent expression levels of each gene.

### Sample preparation for western blot analysis

The samples used for nuclei enrichment were obtained from primordia at stages 3–4 of the WT and cΔ*gcd1* mutant (expressing the GCD1-3xFlag protein). Caps were collected by dissection, as *gcd1* is highly expressed in the cap. Caps were pulverized in liquid nitrogen using mortar and pestle and further homogenized in a nuclei isolation buffer (NIB, 50 mM PIPES/KOH pH 7.0, 1 M hexylene glycol, 10 mM MgCl_2_, 0.2% Triton X-100, 1 mM DTT, 1 mM PMSF, 1x protease inhibitor cocktail (cOmplete, EDTA-free, Merck Millipore) and 25 µM MG132). The homogenate was centrifuged at 200 x *g* for 2 min at 4°C. The pellet was re-suspended in NIB, and centrifuged at 200 x *g* for 2 min at 4°C. This step was repeated. The sample was then filtered through a 40 µm cell strainer (Avantor), and the filtrate was centrifuged at 200 x *g* for 2 min at 4°C. The resulting supernatant (served as the total extract) was further centrifuged at 1,200 x *g* for 10 min at 4°C to separate the cytoplasmic fraction (supernatant) from the enriched nuclear fraction (pellet). Samples were supplemented with Laemmli sample buffer, boiled for 5 min, centrifuged for 5 min at 17,000 x *g* at room temperature and subjected to SDS-PAGE followed by semi-dry transfer to PVDF (Merck Millipore) and western blotting using the following antibodies. Primary antibodies: anti-FlagM2 (1:10,000, Sigma), anti-Histone H3 (1:5,000, Abcam) and anti-αTubulin (1:10,000, Sigma). Secondary antibodies: Goat anti-mouse IgG-HRP (1:10,000, Dako) and Goat anti-rabbit IgG-HRP (1:10,000, Dako).

### Phylogenetic analysis of *gcd1*

For the phylogenetic analysis of *gcd1*, we selected 189 published species^8^ across Agaricomycotina (168 species), Pucciniomycotina (seven species), Ustilaginomycotina (six species) and Ascomycota (seven species from Pezizomycotina, Taphrinomycotina, and Saccharomycotina) (G189 dataset). The identification of the *gcd1* ortholog genes was based on a previous publication^8^. The GCD1 homologs were aligned by mafft (version 7)^71^. Phylogenetic analysis was carried out in RAxML (version 8.2.12) under the CAT + WAG model with 500 rapid bootstrap replicates^72^.

### Copy number examination of proteins from Con7 subfamily and GH30_3 family

To assess the conservation of GCD1, a protein similarity search was conducted using the GCD1 protein sequence (CopciAB_374546) as a query against a dataset of 655 fungal protein sequences (G655 dataset) derived from the 655 fungal genomes which were downloaded from JGI and NCBI before October 22, 2024 (Supplementary Table 9), and the recent copy numbers of GCD1 was visualized in the corresponding species tree. To do this, an MMseqs2 was run in ultrasensitive mode (-s 7.5) to detect distant homologs. Significant hits were defined using an E-value threshold of 1e-5 (-e 1e-5), retaining up to 1,000,000 target sequences per query (--max-seqs 1000000). To confirm that the 741 initial hits belong to the same gene family, a second MMseqs2 search was performed using the 741 proteins as queries against the entire G655 dataset. Considering only the best hit per target species, 732 proteins showed at least 97% overlap in their target gene sets with the original group members of the first MMseqs2 search. After manual curation of ambiguous cases, 730 proteins were ultimately retained as members of the GCD1 gene family.

For species tree inference, we used the 277 marker genes identified by Szánthó et al. (2025)^73^as the starting point. Reciprocal Best Hit (RBH) searches were conducted between the CopciAB genome (query) and the G655 dataset (target) to identify single copy orthologs. Multiple sequence alignments were performed using the MAFFT L-INS-i algorithm and trimmed using TrimAl v1.2 with the strict option. Alignments shorter than 150 amino acids or containing fewer than 500 species were discarded, resulting in 221 single-copy clusters comprising 74,376 aligned sites used for tree reconstruction. Phylogenetic inference was performed using IQ-TREE v1.6.12 with the LG+G substitution model and 1,000 ultrafast bootstrap replicates. A constrain was applied for the tree reconstruction with this topology (((((((((Mixos1, Neoirr1), Mucci2), (Basme2finSC, Entmu)), Olpbor1), Allma1), Ganpr1), Partr), Rozal1_1), Nter) based on former topologies^25^.

The conservation of proteins from GH30_3 family was identified in a published dataset^34^ based on the protein ID of *C*. cinerea (CopciAB_381637), and the species tree and gene family evolution analyses were obtained from that source.

### Harvesting, fixation and embedding of the primordia

The harvesting process involved excising an agar cube bearing hyphal knots or primordia, and subsequently transferring it to a Half-strength Karnovsky’s Fixative^74^ consisting of 1% paraformaldehyde and 2.5% glutaraldehyde. Samples were incubated overnight at 4°C with rotational agitation. For embedding, the samples were dehydrated by incubating in an ascending ethanol series with 10%, 30%, 50%, 70%, 90% and finally 100 % ethanol, respectively with rotational agitation at room temperature. Following this, the samples were embedded in resin TV7100 (Hareus Kulzer, Wehrheim, Germany). The block was sliced into 10 µm thick sections by a rotary microtome (Leica HistoCore Autocut R) equipped with a tungsten carbide knife (16 cm, profile-d). The sections were stained by 0.01% w/v methylene blue (Sigma-Aldrich) and analyzed using a Leica DM 2500 Microscope with a camera attached.

## Supplementary Figures

**Supplementary Fig. 1.**
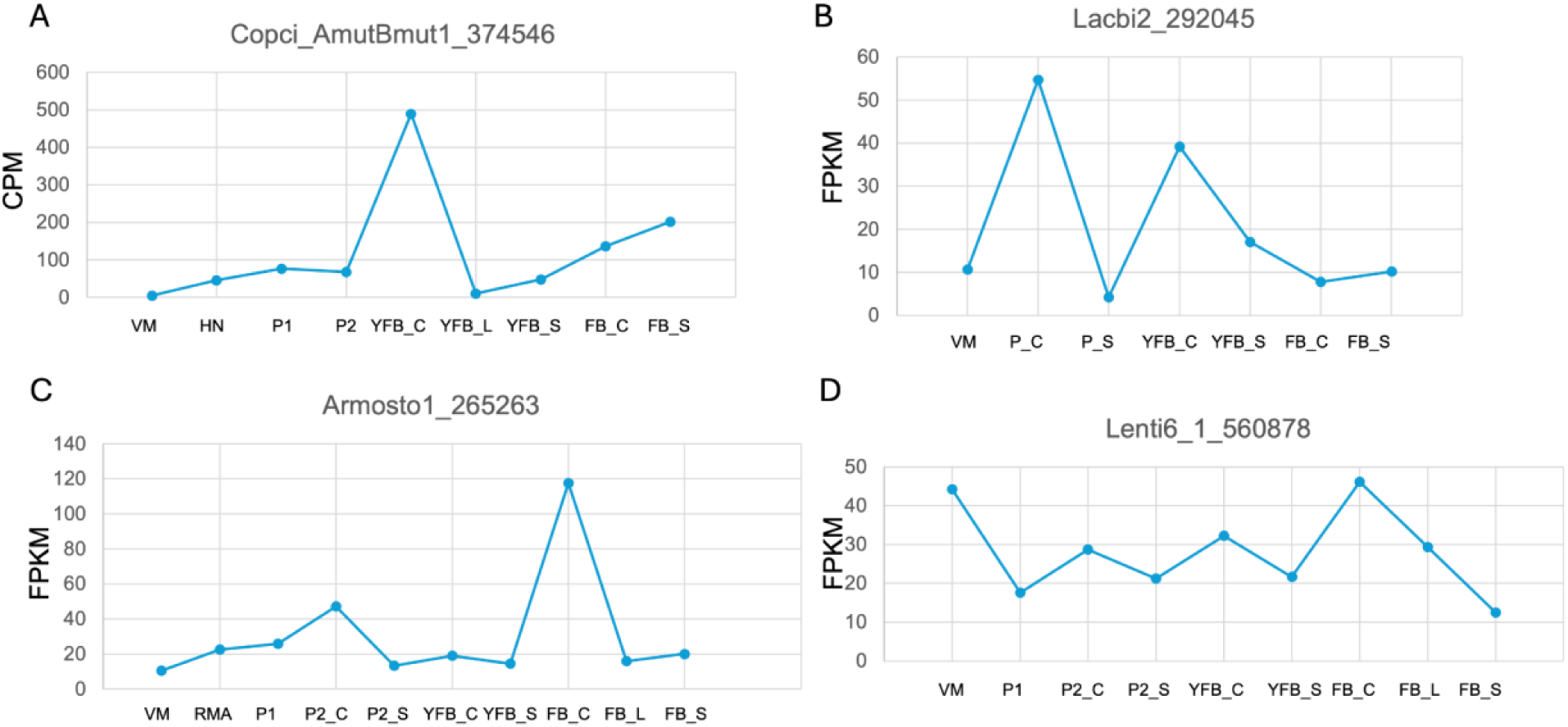
Expression levels of *gcd1* in different mushroom-forming fungi. **A.** The expression level of *gcd1* in *C. cinerea* across developmental stages^15^. VM: vegetative mycelium, HN: hyphal knot, P1: primordium stage 1, P2: primordium stage 2, YFB_C: young fruiting body cap, YFB_L: young fruiting body lamella, YFB_S: young fruiting body stipe, FB_C: mature fruiting body cap and lamella, FB_S: mature fruiting body stipe. **B.** The expression level of *gcd1* ortholog in *Laccaria bicolor* (Lacbi2_292045)^16^ across developmental stages: vegetative mycelium, P_C: primordium cap, P_S: primodium stipe, YFB_C: young fruiting body whole cap, YFB_S: young fruiting body stipe. FB_cap: fruiting body whole cap, FB_stipe: fruiting body stipe. **C**. The expression level of *gcd1* ortholog in *Armillaria ostoyae* (Armosto1_265263)^17^ during the different developmental stages. VM: vegetative mycelium, RMA: rhizomorphs, P1: primordia stage 1, P2_C: primordium stage 2 cap, P2_S: primordium stage 2 stipe, YFB_C: young fruiting body whole cap, YFB_S: young fruiting body stipe, FB_C: fruiting body cap, FB_L, fruiting body lamella, FB_S: fruiting body stipe. **D**, The expression level of *gcd1* ortholog in *Lentinus tigrinus* (Lenti6_1_560878)^18^ across developmental stages. VM: vegetative mycelium, P1: primordium stage 1, P2_C: primordium stage 2 cap, P2_S: primordium stage 2 stipe, YFB_C: whole young fruiting body cap, YFB_S: young fruiting body stipe, FB_C: fruiting body cap, FB_L: fruiting body lamella, FB_S: fruiting body stipe.

**Supplementary Fig. 2.**
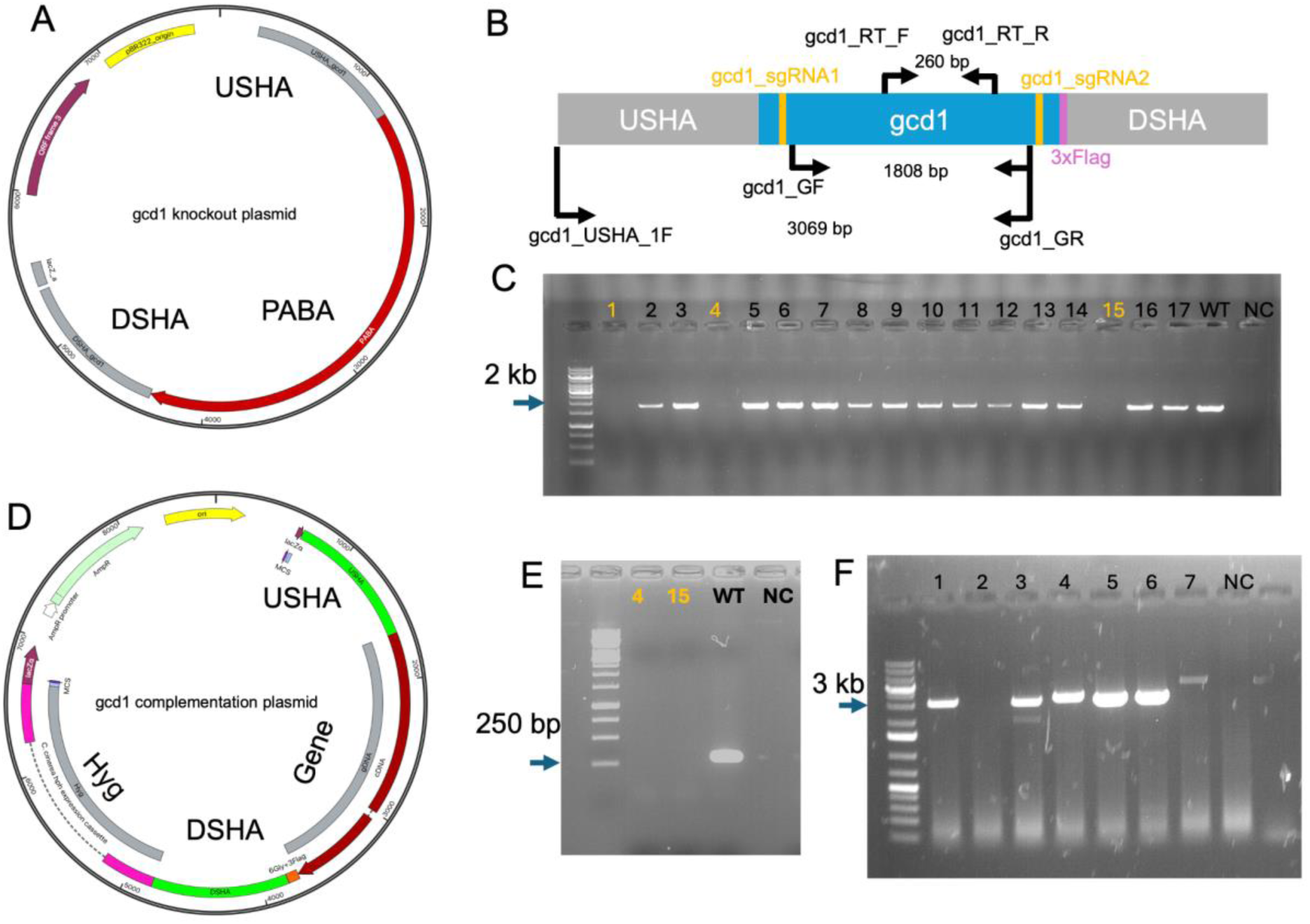
The plasmid map and screening result of the *gcd1* deletion and complementation. A, the map of the plasmid used for generating knock-out mutants. USHA and DSHA represent the upstream and downstream homology arms (1000 bp), respectively; PABA represents the para-aminobenzoic acid gene fragment. B, Schematic representation of the *gcd1* gene indicating the positions of screening primers, Flag tag, and single guide RNAs (sgRNAs). C, PCR screening of the *gcd1* knock-out candidate strains using gel electrophoresis using the gcd1_GF/gcd1_GR primer pair, the positive transformants are marked as orange, WT represents positive control by using the gDNA template from the WT strain, NC represents no template control. D, the map of the plasmid used for complementation. USHA and DSHA represent the upstream and downstream homology arms (1000 bp), respectively; Hyg represents the hygromycin B phosphotransferase gene fragment. E, RT-PCR testing for knocking out strain using the gel electrophoresis (marked as orange) using the gcd1_RT_1F/gcd1_RT_1R primer pair, and cDNA as the template. WT represents positive control by using the cDNA template from the WT strain, NC represents no template control. F, PCR screening of the complementation strains using the gcd1_USHA_1F/gcd1_GR primer pair. Those samples were selected as complementation strains from which fragments of the appropriate size (3069 bp) could be amplified from the genomic DNA.

**Supplementary Fig. 3.**
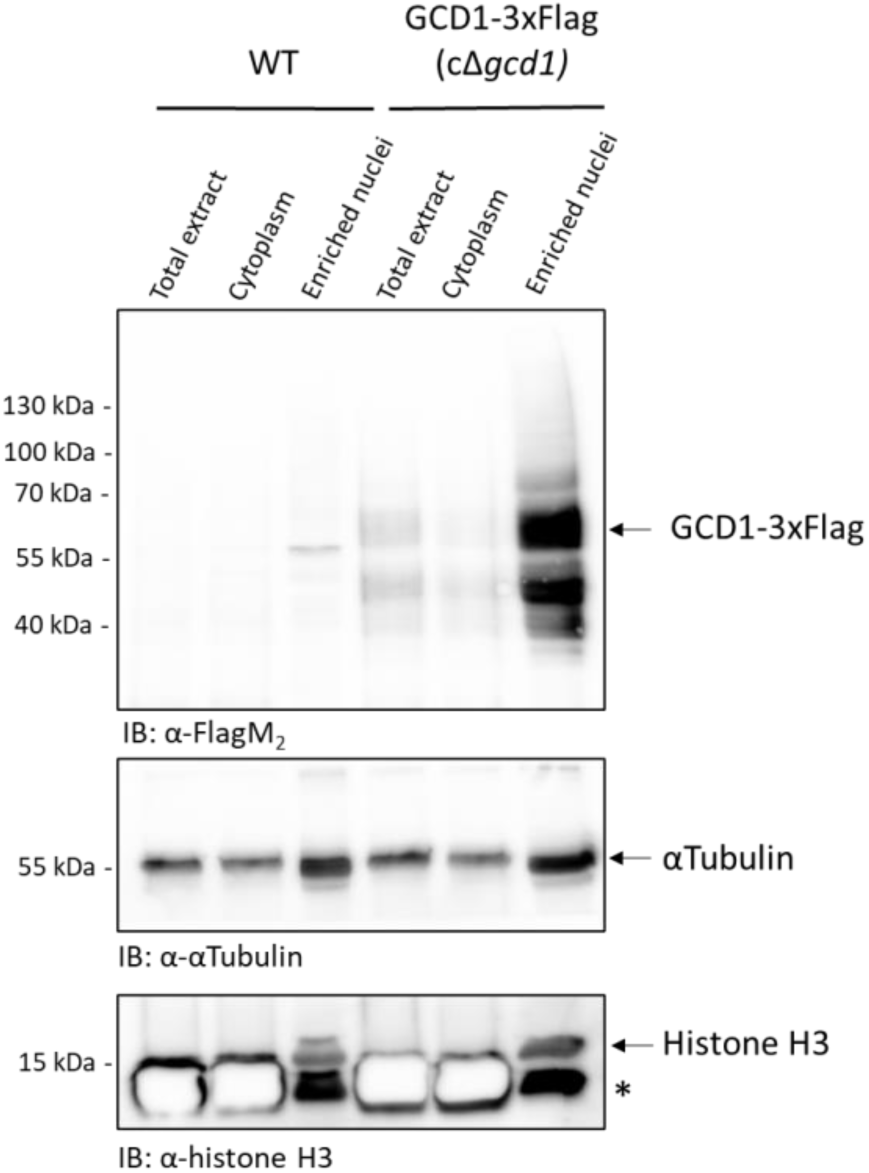
GCD1-3xFlag is highly enriched in the nuclei prepared from the cap of the complemented cΔ*gcd1* mutant. Total extracts, cytoplasmic fractions and enriched nuclei prepared from WT or cΔ*gcd1* mutant caps were subjected to SDS-PAGE and analysed by western blotting using the indicated antibodies. As shown, GCD1-3xFlag was detected in the total extract and was highly enriched in the nuclear fraction of the complemented cΔ*gcd1* mutant, while it was not detectable in the WT samples. αTubulin served as a loading control, and Histone H3 served as a nuclear marker. Asterisk (*) indicates an aspecific band. IB: immunoblot.

**Supplementary Fig. 4.**
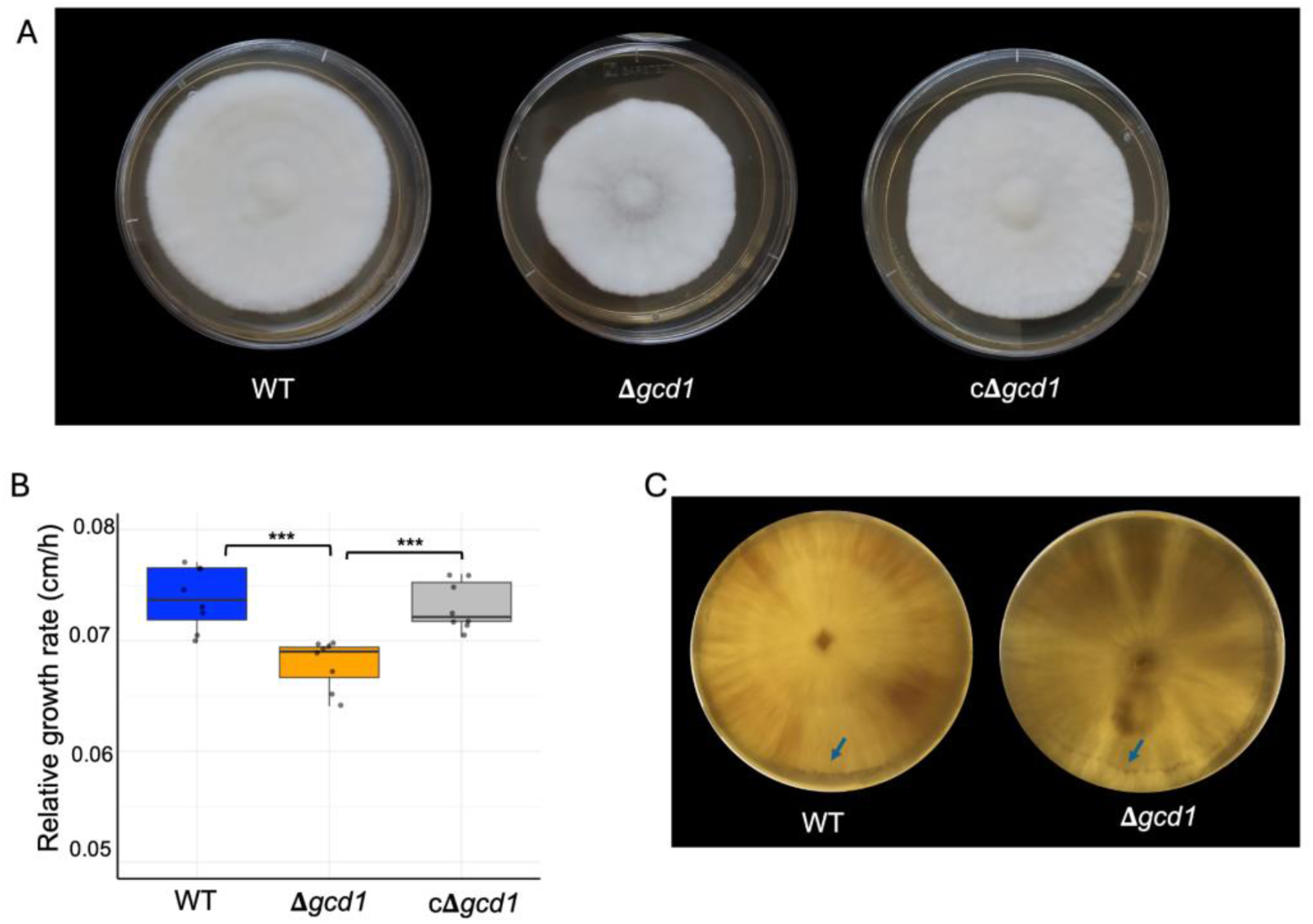
Mycelium growth and hyphal knot forming of the WT and Δ*gcd1* strains. A, mycelial growth of WT, Δ*gcd1* and cΔ*gcd1* strains on the YMG medium for 5 days after inoculation. B, boxplot showing the vegetative mycelium growth rate among WT, Δ*gcd1*, and cΔ*gcd1* strains. Eight replicates were used for each strain, the Student’s t test was used to compare the means between each group, *** p < 0.001 (two-way ANOVA). C, the hyphal knot ring (arrows indicated) formed after 2 hours of light induction followed by 24 hours of incubation in the dark at 28°C.

**Supplementary Fig. 5.**
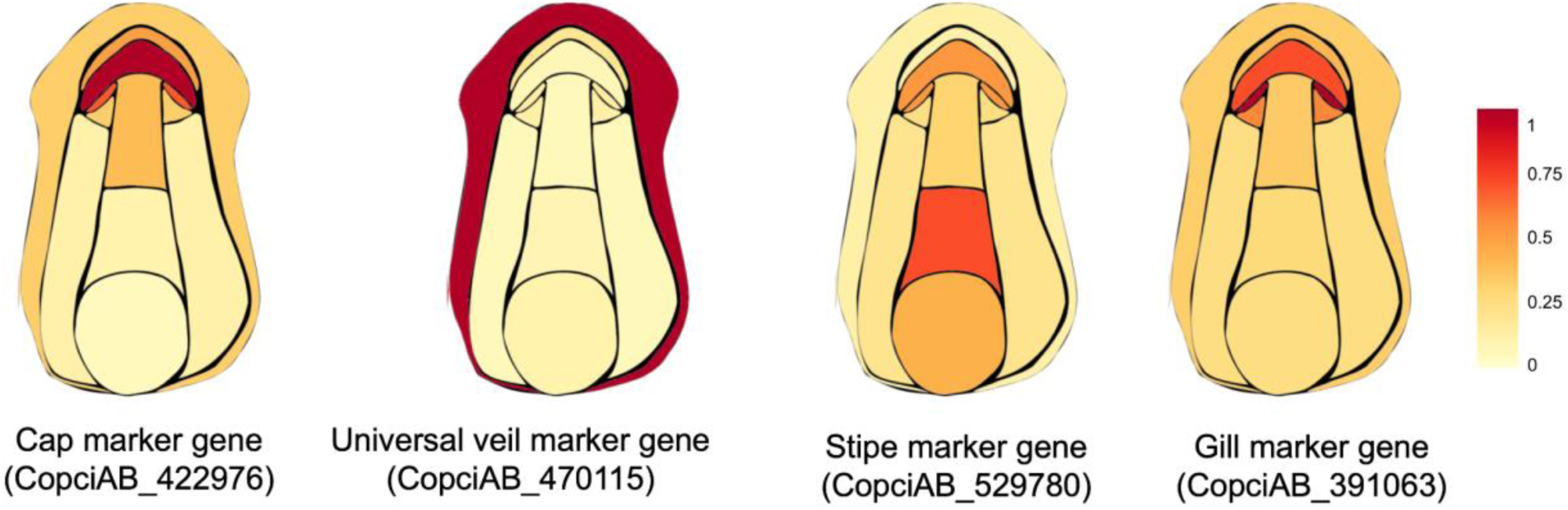
Expression image heatmap of tissue marker genes. Log2-transformed normalized counts were used for the heatmap, and the color scale was normalized from 0 to 1 to represent expression levels of each gene.

**Supplementary Fig. 6.**
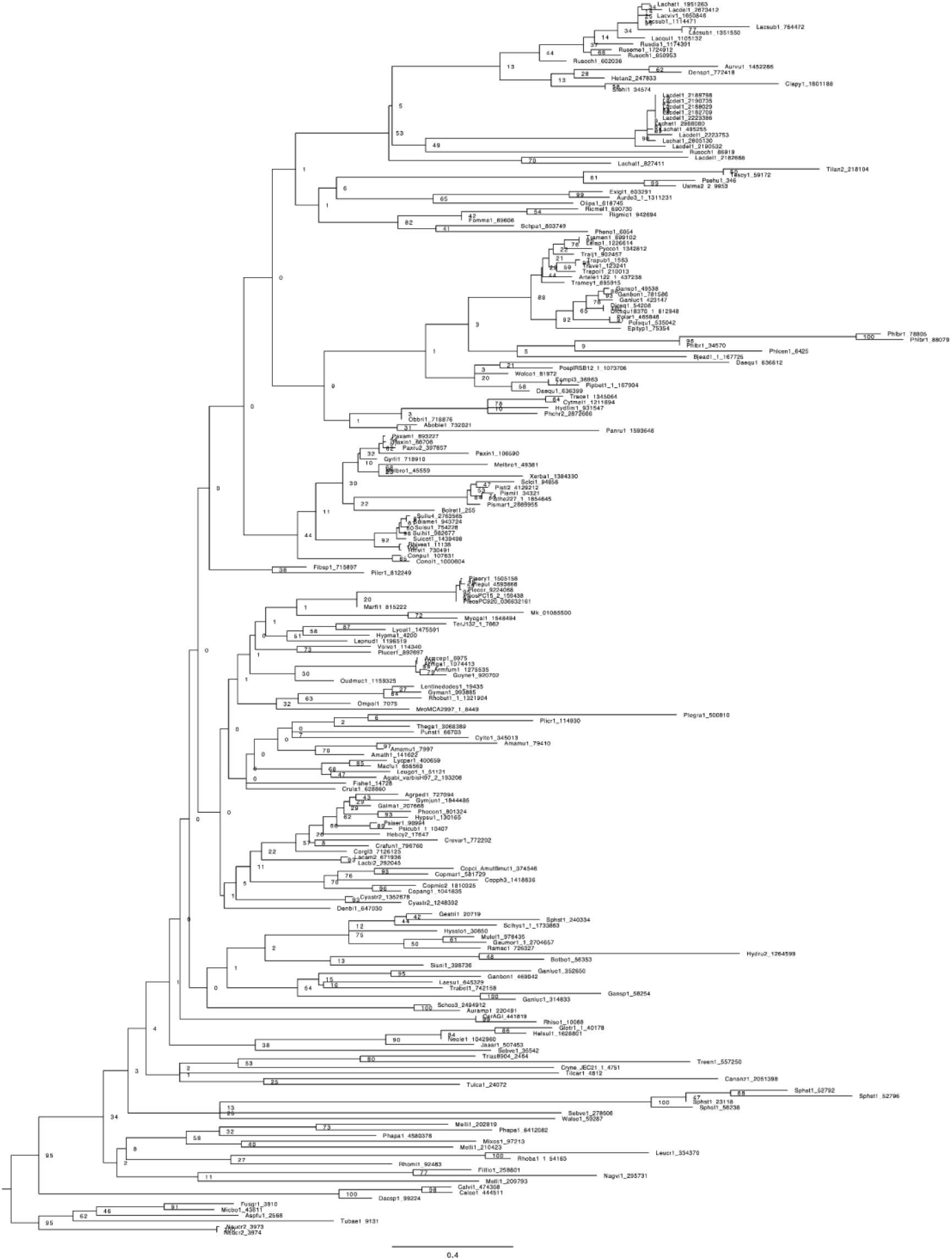
Phylogenetic analysis by using GCD1 homologs identified in 189 published genomes.

**Supplementary Fig. 7.**
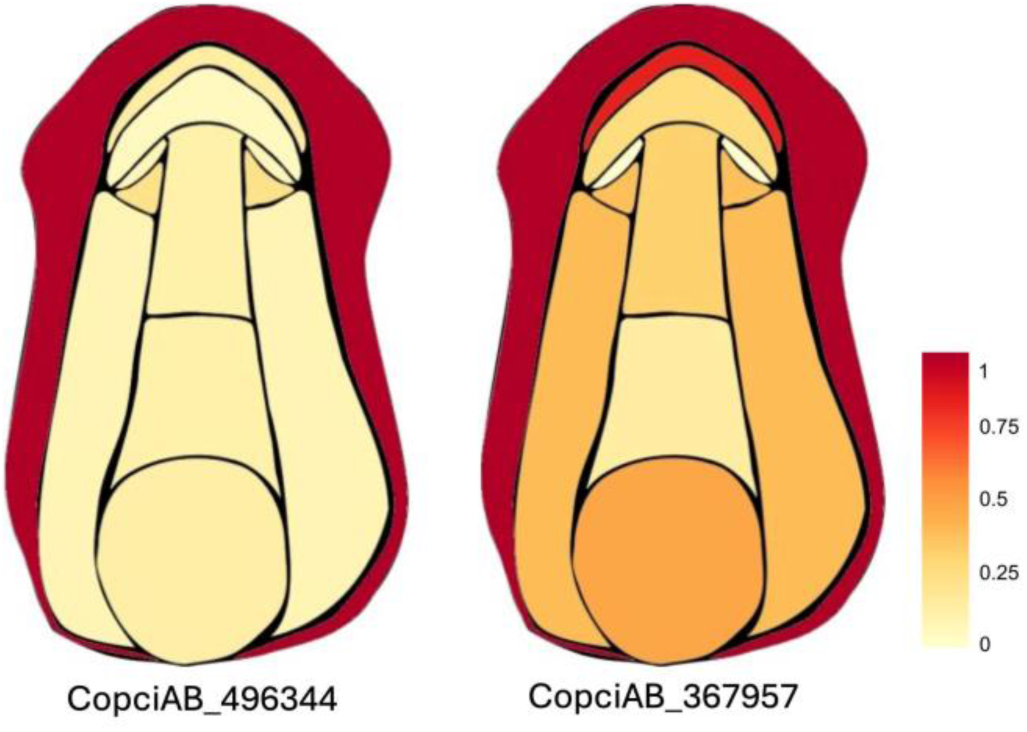
Expression image heatmap of two defense-related genes. The Log2-transformed normalized counts were used for the heatmap, and the color scale was normalized from 0 to 1 to represent expression levels of each gene for each tissue type.

