## Supplementary tables for "Functional characterization of a Con7-related transcription factor in *Coprinopsis cinerea* indicates evolutionary conservation of morphogenetic roles": Table 1.docx

Table 1 Custom gene sets used for enrichment analysis

| Terms shown in the figure | Description | References |
| --- | --- | --- |
| YFB_cap_specific | Cap specific gene identified in young fruiting body (FC > 2) | Krizsán et al., 2019 |
| YFB_gill_specific | Gill specific gene identified in young fruiting body (FC >2) | Krizsán et al., 2019 |
| YFB _stipe_ specific | Stipe specific gene identified in young fruiting body (FC > 2) | Krizsán et al., 2019 |
| FB-initiation | Gene expression increased at least fourfold in hyphal knots and stage 1 primordia, relative to vegetative mycelium | Krizsán et al., 2019 |
| Core starvation response | Genes induced by designated starvation condition | Hegedüs et al., 2025 |
| Late light response | Genes induced within two hours post light exposure | Hegedüs et al., 2025 |
| Early light response | Genes induced after 6 hours post light exposure | Hegedüs et al., 2025 |
| Aerial mycelium specific | Aerial mycelium specific gene identified without starvation | Hegedüs et al., 2025 |
| Attached mycelium specific | Attached mycelium specific gene identified without starvation | Hegedüs et al., 2025 |
